# Time-resolved neural and experience dynamics of medium- and high-dose DMT

**DOI:** 10.1101/2024.12.19.629418

**Authors:** Evan Lewis-Healey, Carla Pallavicini, Federico Cavanna, Tomas D’Amelio, Laura Alethia De La Fuente, Debora Copa, Stephanie Müller, Nicolas Bruno, Enzo Tagliazucchi, Tristan Bekinschtein

## Abstract

N,N-Dimethyltryptamine (DMT) is a fast-acting psychedelic drug that induces a radical reorganisation of conscious contents and brain dynamics. However, our understanding of how brain dynamics support psychedelic-induced conscious states remains unclear. We therefore present a repeated-measures dose-dependent study of the subjective and neural dynamics induced through DMT under naturalistic conditions. Nineteen participants received either a 20mg or 40mg dose of freebase DMT across two sessions in a blinded, counterbalanced order. Electroencephalography (EEG) data and time-resolved measures of subjective experience (Temporal Experience Tracing) were collected. Both doses of DMT induced rapid changes in experience dimensions, with the 40mg dose inducing more extreme visual hallucinations and emotionally intense experiences. Strikingly, lempel-ziv complexity, previously hailed as a robust phenomenological correlate within the psychedelic-state, was the least strongly associated neural marker. These findings suggest that the relationship between neural complexity and phenomenology in psychedelic states is less clear than originally hypothesised.

---

Much like lesion studies heralded a profound shift in the scientific understanding of the brain [1], psychedelic drugs are increasingly hailed as promising pharmacological candidates to advance the neuroscience of consciousness [2, 3]. They present a unique opportunity to radically and reversibly perturb both the structure of experience and neural dynamics, thereby facilitating mapping the relationship between the mind and brain, one of the fundamental goals of the neuroscience of consciousness.

Among the class of serotonergic psychedelics - agonists at the 5-HT_2A_ receptor - is N,N-Dimethyltryptamine (DMT). When administered intravenously or inhaled in freebase form, DMT can induce brief yet profound alterations to subjective experience [4–10]. Previous research has found that when the dose of DMT exceeds a certain threshold, participants enter a highly immersive state, colloquially known as “breaking through”, comprising the dissolution of space and time, distortions in the sense of self, and the perception of intricate and complex geometric hallucinations with eyes closed [4, 5]. Further, a large proportion of DMT users report perceiving “entities” within this state, described as sentient otherworldly beings who are often attempting to convey information to the user [11]. At the neural level, there are a multitude of deviations from normal waking consciousness associated with both DMT administration and serotonergic psychedelics more broadly. These include decreases in global alpha power [6, 9, 10, 12–14], broadband spectral changes [12, 15], and increased global functional connectivity [10, 16, 17]. Additionally, an overarching theory within the neuroscience of psychedelics (the Entropic Brain Theory [18, 19]) posits that neural entropy is associated with the ‘richness’ of subjective experience. This is supported by studies finding that Lempel-Ziv (LZ) complexity increases as a function of the phenomenologically rich psychedelic state [9, 10, 20, 21], and decreases in states of consciousness that lack subjective experience [22–24]. Moreover, previous research has found that neural complexity can track the subjective intensity of the psychedelic experience [9, 20, 21], further supporting the basic tenet of the entropic brain theory.

However, one fundamental issue emerges when attempting to measure subjective experiences: discrete Likert scale methodologies fail to capture the inherent dynamic complexity of phenomenology [10, 25]. This is a pertinent issue within psychedelic neuroscience as the psychedelic state has nuanced and idiosyncratic dynamics [26], with subjective experiences undulating with the pharmacodynamics of the substance. It is therefore of utmost importance to integrate more complex measures of phenomenology with neuroimaging techniques, dubbed neurophenomenology [2, 27, 28], to more successfully bridge the gap between experience and brain dynamics under psychedelics.

The present study addresses this gap in research concerning the failure to apply complex and quantitative models of phenomenology to the dynamic experiences induced by DMT. We build on previous research that have sought to integrate time into their neurophenomenological analysis of psychedelics [9] by using Temporal Experience Tracing (TET; [29–32]). TET is a phenomenological methodology whereby participants retrospectively trace the subjective intensity of experience dimensions in time, yielding continuous data (see Methods), rather than discrete data points as generated by Likert scales. By doing so, TET permits the production of multidimensional phenomenological timeseries - temporally-resolved reproductions of experience which can be more easily and fruitfully paired with objective brain-imaging dynamics. We use TET in combination with electroencephalography (EEG) in a within-participants dose-dependent study of DMT under naturalistic conditions. These naturalistic conditions allowed participants to experience the effects of DMT in a relaxed and familiar setting, favouring the recording of high-quality data with minimal artifacts associated with stress and anxiety [6]. Due to the broad swathe of neural changes associated with the psychedelic state, we compute a variety of neural features that fall into overarching families, and associate these to continuous subjective experience dimensions (see Methods). We broadly hypothesised that a) 40mg doses would be significantly higher in intensities for all dimensions, and b) neurophenomenological associations would be most strong amongst information theory measures, oscillatory alpha, and oscillatory delta power.

## Results

We employed a repeated-measures within-subjects blinded 2×2 design under naturalistic conditions, with dose (20mg and 40mg of freebase DMT) as a counterbalanced between-session factor and condition (DMT and resting-state (RS)) as a within-session factor (Figure 1A). Nineteen participants (17 males; 34.6±5 years; 4.7±5 previous DMT experiences) were included in the study, with subjective experience dynamics of each condition measured through TET (Figure 1B), and neural dynamics of each condition measured through EEG, totalling 76 recordings (19 sessions x two conditions x two doses). After preprocessing, two EEG sessions were excluded from the DMT condition (one due to excessive slow-wave activity and one due to to more than 50% of the epochs rejected) and one from the RS condition (due to excessive slow-wave activity). The application in which we collected TET dimensions also failed to load for one participant for the DMT condition, and was therefore unavailable for subsequent analysis.

**Figure 1:**
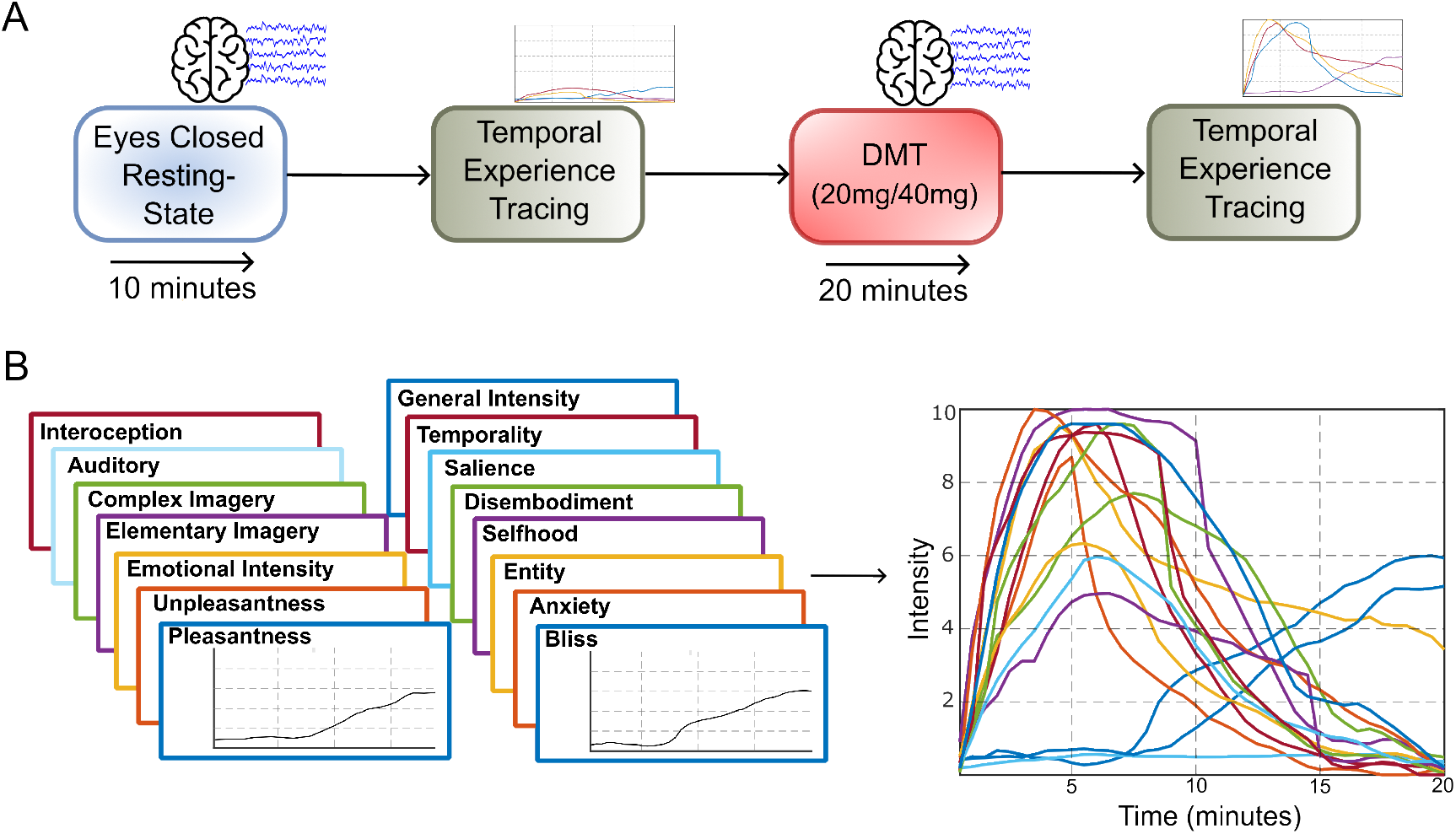
A schematic of the experimental design and phenomenological methodology (Temporal Experience Tracing; TET) used within this study. **(A)** The format of the dosing-day session each participant underwent (participants attended two dosing-day sessions). In the first section, resting-state (RS) EEG activity was collected, whereby the participants were instructed to maintain wakefulness with eyes closed. Succeeding this, participants traced the retrospective subjective intensity of 15 experience dimensions for the 10-minutes of the RS condition. After this, when the participant was ready, they received either 20mg or 40mg of freebase DMT (the order in which they received this was counterbalanced and randomised). Once fully smoked, the participant lay back in a supine position, while EEG activity was recorded. Over the duration of the 20-minutes, a recording of bells at two-minute intervals were played to serve as temporal markers. Once the 20-minutes had finished, and once the participant was ready, they then completed the retrospective traces of the 15 dimensions of subjective experience. **(B)** The 15 dimensions used within this study (left) were concatenated into vectors to form a multidimensional matrix of subjective experience for the RS and DMT conditions. The panel on the right is data from the 40mg dose of S01.

After completing the TET dimensions for each DMT condition, participants guessed which dose they thought they had received; 26 out of the 38 sessions were guessed correctly (14/19 (74%) correct in the 40mg session, 12/19 (63%) correct in the 20mg session). A chi-square test of independence revealed no significant differences between the blinding of the two conditions χ^2^ (1, N = 38) = 0.49, *p* = .49. However, a chi-square goodness of fit test revealed that participants correctly guessed the condition above chance across both dose conditions χ^2^ (1, N = 38) = 5.16, *p* = .02.

### DMT doses induce nuanced experience dynamics across phenomenological dimensions

We took a dual approach, first ignoring the temporal aspect of the TET methodology, and second, integrating it within our analysis to utilise its full potential. The results, as expected, provide phenomenological nuance when comparing between doses. Our first analysis entailed statistically comparing all TET data generated from each dose condition (40 data points per dose per participant) in a mixed modelling fashion, akin to Likert scale analyses but with more data to power the analysis. We computed a linear mixed model for every phenomenological dimension using only TET data for each DMT dose, factoring subject and session as nested random effects within the model. This approach disregards the temporal aspect of phenomenology, simply investigating the average differences in dimension intensity between doses. Using this approach, Elementary Imagery (−0.581, *t*(1,478) = -3.69, *p* = 0.003), General Intensity (−0.58, *t*(1,478) = -3.18, *p* = 0.008), Complex Imagery (−0.672, *t*(1,478) = -3.16, *p* = 0.008), and Selfhood (−0.832, *t*(1,478) = -3.07, *p* = 0.008) were significantly lower in the 20mg DMT condition compared to the 40mg condition (Supplementary Figure S1). However, no other phenomenological dimensions were significantly different after FDR correction.

Following this, and to use the full richness of the dynamics of TET, we included time within each linear mixed model, comparing each phenomenological timestep to the average restingstate data point of that specific phenomenological dimension (see Methods). For example, for the dimension Elementary Imagery, the formula was ‘Elementary Imagery Time*Dose + (1| Subject:Session)’, with ∼ ‘Time’ and ‘Dose’ as categorical variables. A summary of the results can be found in Figure 2A. Broadly, through the incorporation of time within our statistical analysis, we found Emotional Intensity, Salience, Selfhood and Interoception to be most significantly different to the restingstate average for the longest period of time across both DMT dose conditions (the strength of the main effect coefficients can be found in Supplementary Figure S2). Further to this, dose interaction effects revealed that the 40mg dose yielded stronger differences between the RS average across most dimensions for at least three minutes of the session (Selfhood (16 minutes); Complex Imagery (13 minutes); Elementary Imagery (12 minutes); General Intensity (9.5 minutes); Salience (9 minutes); Unpleasantness (9 minutes); Emotional Intensity (5 minutes); Entity (5 minutes); Temporality (4 minutes); Anxiety (3 minutes). There were no interaction effects for the dimensions Interoception, Pleasantness, Auditory, Disembodiment, and Bliss.

**Figure 2:**
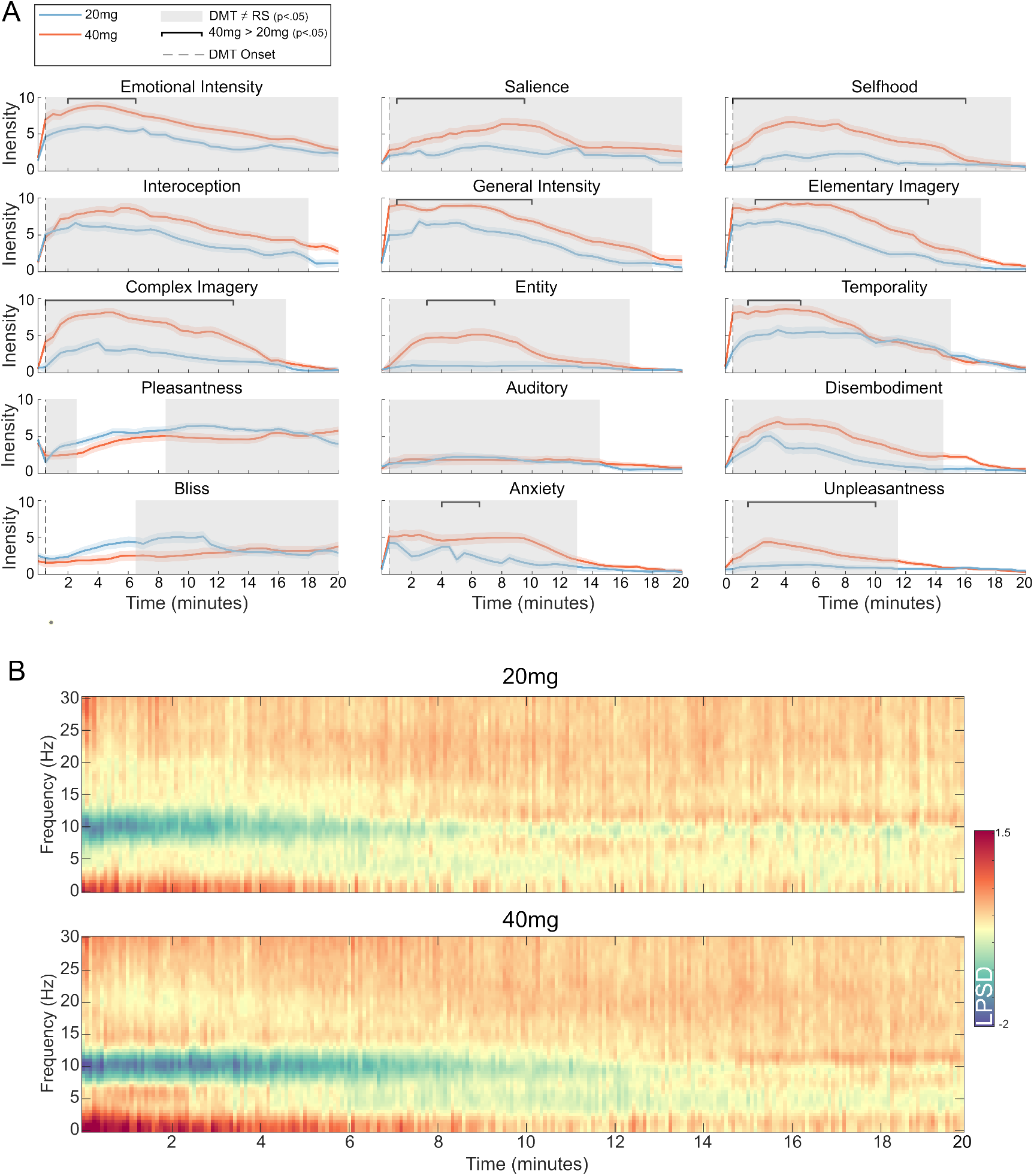
Phenomenological reports and EEG spectral power present distinct dynamics that distinguish between DMT doses. **(A)** Time series plots of the 40mg (red) and 20mg (blue) dose for each phenomenological dimension. Standard error of the mean is shaded. The first time-point is the average for the whole duration of the resting-state (RS) condition, with grey dashed lines aligned to the first phenomenological data point from the DMT condition. Grey boxes indicate that the DMT condition, regardless of dose, was significantly different from the RS average for that dimension at that time point. Black bars indicate a dose-interaction effect, whereby the difference between resting-state and DMT was moderated by a dose effect (the 40mg dose was higher than the 20mg dose in all significant interaction effects). Dimensions are ordered by the strength of main effect coefficient sizes (Supplementary Figure S2). **(B)** Average logarithmic power spectral density (LPSD) changes for 20mg (top) and 40mg (bottom) doses subtracted from RS LPSD. Statistical analyses were not computed here, as our primary aim was to allow the phenomenology to guide our statistical analyses. Neurophenomenological analyses can be found in Figure 4.

### Oscillatory alpha power and permutation entropy yield strongest neurophenomenological associations

We also conducted neurophenomenological analyses to identify how subjective experience dynamics related to EEG dynamics during the DMT-state. We computed a variety of neural features broadly associated with the psychedelic-state, using a similar approach to previous research[33, 34]. However, within this analysis, we focused on how neural features related to phenomenology of the DMT-state, rather than statistically analysing the difference between RS and psychedelic-state conditions. Broadly, the neural features computed were categorised into three families of markers: information theory, spectral, and connectivity (Figure 3A, 3B, and 3C respectively). We elaborate on each measure and its computation in the Methods. Co- variance matrices between each (global) neural feature can also be found in Supplementary Figure S3. It is of note that each neural feature was standardised to have mean of 0 and variance of 1. Therefore the coefficients are comparable across neural features, and convey the strength of association between subjective experience dimensions and neural features, similar to effect sizes [35].

**Figure 3:**
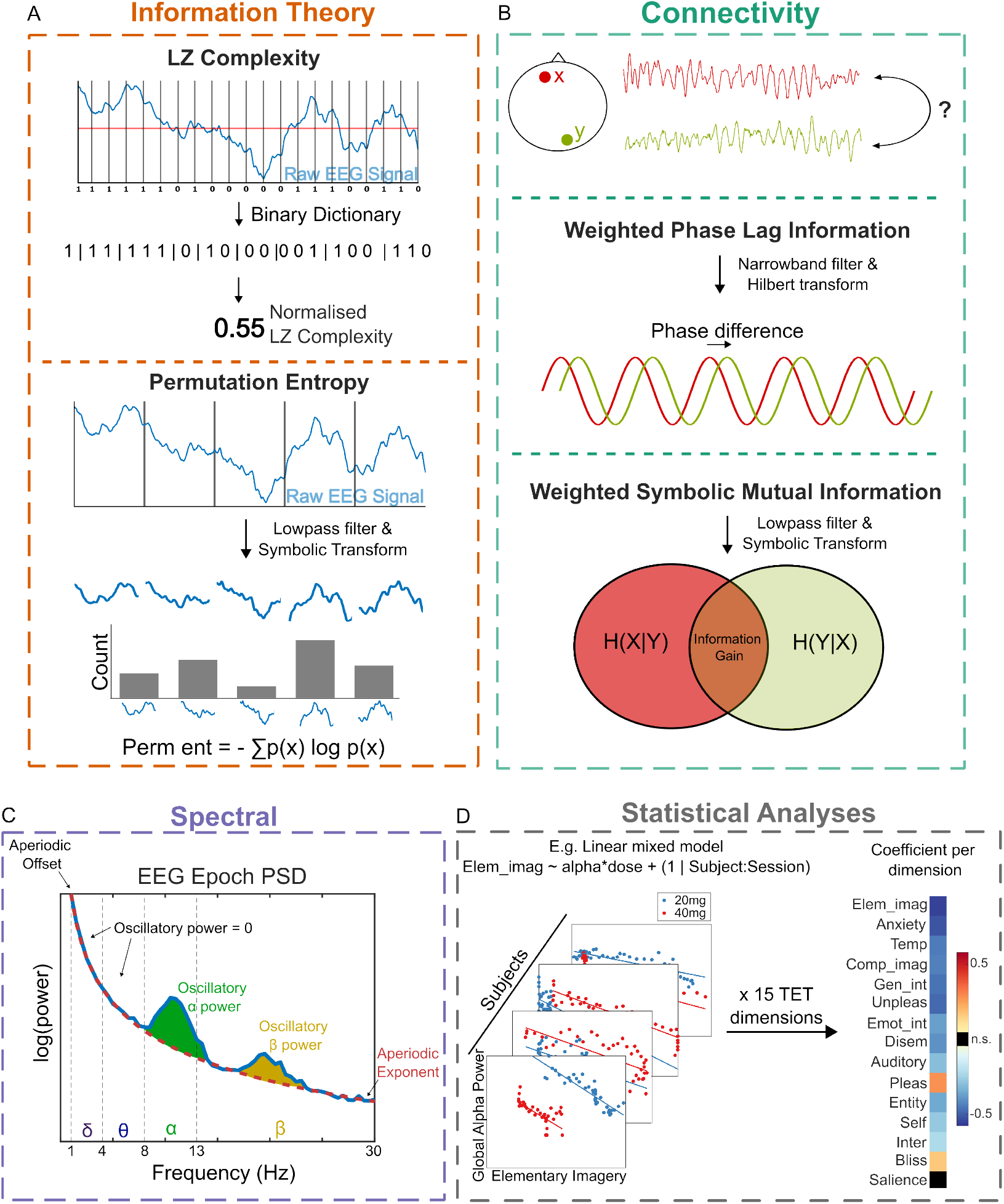
A schematic illustration of neural feature computation, and how features are statistically associated with TET dimensions. **(A)** Within the information theory family, Lempel-Ziv complexity (LZ) and Permutation Entropy were computed. LZ (above) involves binarising the raw EEG signal and evaluating the compressibility of this binarised signal. Permutation entropy (below) is computed through the symbolic transformation of an EEG epoch, which then contributes to a discrete probability distribution of symbols represented for the epoch. Shannon entropy is then computed on this probability distribution, to index the EEG signals level of uncertainty. **(B)** Within the connectivity family, weighted phase lag information (wPLI) and weighted symbolic mutual information (wSMI) were computed to evaluate EEG electrode coupling. wPLI (above) is a linear measure analysing phase-related differences, whereas wSMI (below) is a nonlinear connectivity measure evaluating the information gain of one signal through the observation of another. **(C)** Within the spectral family, we parameterised aperiodic/broadband/arrhythmic and periodic components of the EEG power spectral density (PSD) using the specparam method [36]. The exponent and the offset reflect the aperiodic components, while oscillatory power corresponds to ‘peaks’ of activity above the parameterised exponent, as highlighted in the alpha and beta frequency bands. **(D)** A schematic of the linear mixed modelling approach used. For each phenomenological dimension and each neural feature, we computed a linear mixed model quantifying the overall strength of association. For each neural feature, this yielded 15 coefficients, one for each phenomenological dimension.

To analyse the relationship between TET dimensions and each neural feature, linear mixed models were computed between each phenomenological dimension during the DMT condition and each neural feature, with dose as an interaction effect (Figure 3D).

The results partly corroborated with our hypotheses: oscillatory alpha markers (median effect size = 0.34; std = 0.13)^1^ and permutation entropy markers (median effect size = 0.27; std = 0.12) had the largest coefficients, and were therefore most strongly associated with TET dimensions. However, the majority of the LZ complexity association coefficients (median effect size = 0.04; std = 0.08) were nonsignificant after FDR correction (Figure 4A). Further, when computing power in the canonical frequency bands through the parameterisation of true oscillatory ‘bumps’ in the power spectral density (PSD) after factoring out aperiodic activity (Figure 4B), we found little oscillatory delta power across both DMT and RS conditions (Supplementary Figure S4). When computing the percentage of delta power relative to the spectrum of activity in a conventional manner (see Supplementary Methods), we found that delta power did yield neurophenomenological associations (Supplementary Figure S5 & S6), echoing previous research [9, 10]. However, we contest that this may be more reflective of broadband aperiodic changes in the PSD, rather than an increase in oscillatory delta power [36]. Regarding connectivity markers, wSMI yielded slightly stronger associations (median effect size = 0.15; std = 0.10) than wPLI (median effect size = 0.09; std = 0.07) (Figure 4C). Based on our hypotheses, we also conducted two-sample Kolmogorov-Smirnov tests (due to non-normal distributions and violation of independence of assumptions) between the effect size distributions of specific neural markers, to evaluate whether certain neural markers were more significantly associated with phenomenology. Our targeted analyses revealed that oscillatory alpha power was significantly more associated with TET dimensions than permutation entropy (*D* = 0.69, *p*<0.001), permutation entropy was significantly more associated with TET dimensions than LZ complexity (*D* = 0.25, *p*<0.001), and wSMI was significantly more associated with TET dimensions than wPLI (*D* = 0.76, *p*<0.001). See Figure 4D for distributions of the effect sizes across all neural markers. For all neurophenomenological associations, positive valence dimensions (i.e., Bliss and Pleasantness) yielded the opposite trends to the rest of the TET dimensions. For example, both Pleasantness and Bliss were positively associated with oscillatory alpha power, while the rest of the phenomenological dimensions were negatively associated with oscillatory alpha power. Interestingly, this is in contrast to a previous study with TET and breathwork [31], which found positive and significant associations between both neural LZ complexity and the aperiodic exponent - but not alpha oscillatory power - and the phenomenological dimension of Bliss.

**Figure 4:**
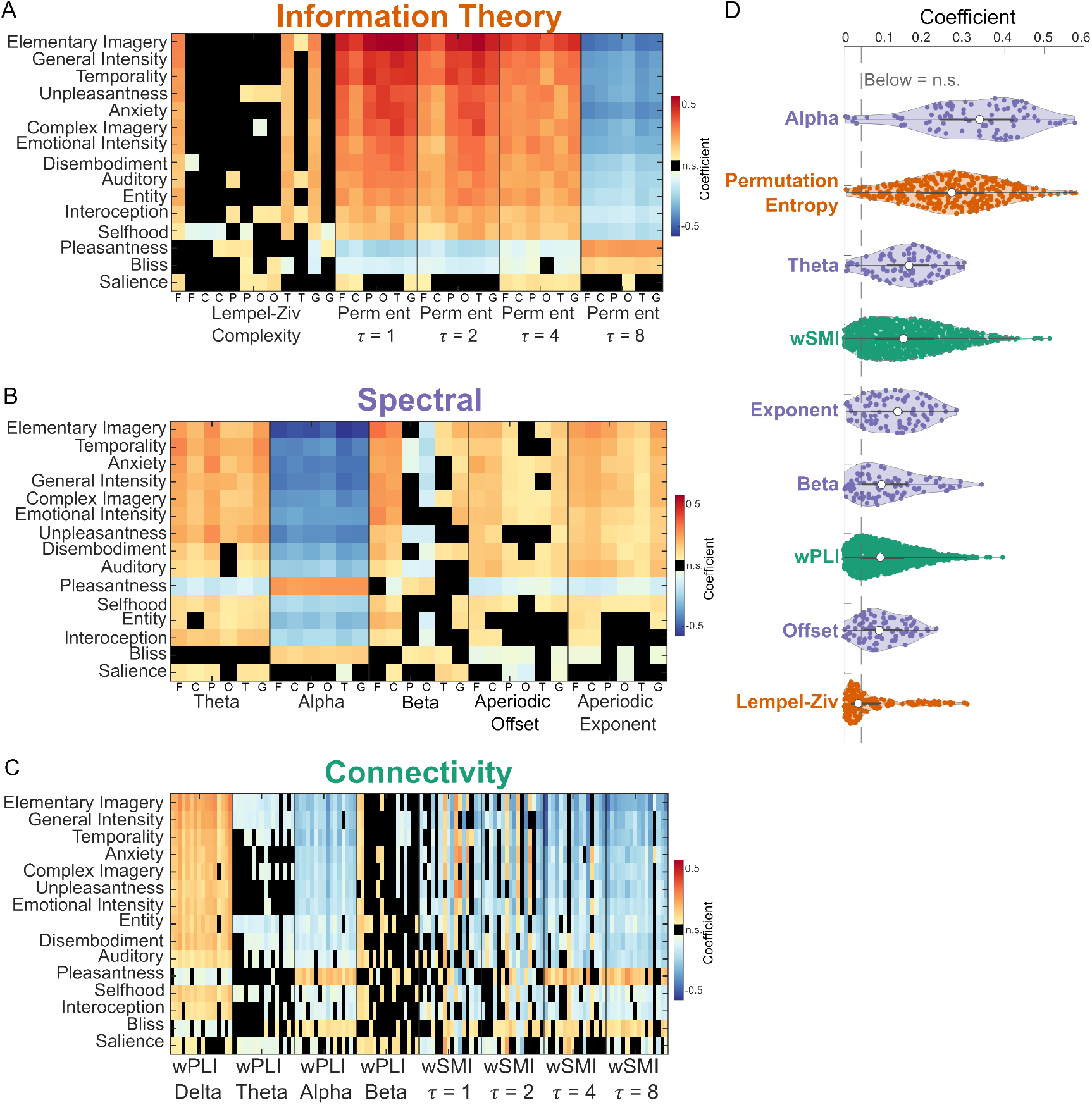
Permutation entropy and oscillatory alpha power most strongly associated with phenomenology during the DMT condition. **(A)** Main effects of the linear mixed model coefficients for the information theory markers. Lempel-ziv complexity is computed through two different means, therefore yielding 12 features (see Methods). Values marked in black were non-significant (*p*>0.05) after FDR correction. F = frontal EEG channels; C = central; P = parietal; O = occipital; T = temporal; G = global. **(B)** Main effects of the linear mixed model coefficients for the spectral markers. Oscillatory delta is not present due to a lack of oscillatory activity in the delta band. **(C)** Main effects of the linear mixed model coefficients for linear (wPLI; weighted phase lag information) and non-linear (wSMI; weighted symbolic mutual information) connectivity markers. **(D)** Distributions of all the coefficient values for each neural feature, ordered by average coefficient size on the y-axis. All coefficients were converted to positive integers for comparison across neural features, with non-significant coefficients also plotted. Median values (white dot) and quartiles (grey whiskers) are in the centre of each violin plot. Any coefficient values below the grey dashed lines were non-significant after FDR correction. The scale of the coefficients are the same for A, B, and C, and phenomenological dimensions are ordered based on the overall strength of association for all neural features within that family matrix.

To complement the main effects of evaluating the relationship between each neural feature and phenomenological dimension across both doses, we also computed interaction effects, assessing whether dose had an impact on neurophenomenological associations (Supplementary Figures S7-S9). When transforming the coefficients to investigate whether the dose interaction strengthened or weakened the association between neural feature and phenomenological dimension, we broadly found that the neurophenomenological associations in the 20mg dose condition were weaker (Supplementary Figure S10). This complements the notion that the experience dynamics between the two doses are distinct, and therefore yield distinct neural dynamics. To validate the assumption that the phenomenological dimensions were meaningfully tracking subjective experiences under the psychedelic state, and may not be more broadly related to phenomenology during normal waking consciousness, we also computed linear mixed models quantifying the degree of association between each TET dimension and each neural feature under the RS condition. We found minimal significant neurophenomenological associations in this condition across all neural features (Supplementary Figure S11).

## Discussion

We studied the neural and subjective experience dynamics of both 20mg and 40mg of inhaled freebase DMT in a repeated-measures within-subjects design under naturalistic conditions. We found that, through the use of TET as a temporally-derived phenomenological methodology, distinct experience dynamics emerged both between both doses (even after similar blinding rates) and across phenomenological dimensions. While DMT broadly induced rapid and radical changes in many dimensions over time, most notably Emotional Intensity, Salience and Selfhood, the high dose was associated with reliably stronger changes, confirming our hypothesis. Conducting neurophenomenological analyses across a range of neural features, we found that experience dimensions were most strongly associated with changes in oscillatory alpha power and permutation entropy, partially conforming our main neurophenomenological hypothesis. Surprisingly, however, LZ complexity, previously hailed as a robust marker of conscious state in a variety of studies [21, 22, 24, 37], was the weakest performing neural feature. Further, the medium dose broadly showed weakened associations between the brain and experience dynamics, and almost no neurophenomenological associations were found when investigating the RS condition. We discuss the implications of these findings below.

While we found reliable differences in only four dimensions when averaging all the data together in a coarse-grained mixed model approach, the TET experience dynamics were nuanced across doses, revealing a rich and more precise phenomenological landscape. This is contrary to our broad hypotheses, which posited that the 40mg dose would yield increased differences to RS phenomenological baselines compared to the 20mg dose across all phenomenological dimensions; some dimensions, such as Interoception, Pleasantness and Bliss were not significantly different at the interaction level for the duration of the session, whereas others, such as General Intensity, Elementary Imagery, and Complex Imagery were for parts of the session, again highlighting dynamically nuanced phenomenological dose-dependencies. One dose-dependent difference of note was valence; for the 40mg dose, retrospective ratings of unpleasantness were reliably higher during minutes 2-10, coupled with transiently higher phases of anxiety. Due to the potential transdiagnostic application of psychedelics for mental health issues [38], and the hypothesis that the subjective experience of psychedelics is necessary to promote positive clinical outcomes [39], these results may be informative for the clinical application of DMT (NCT06094907). As transient negative valence scales with dose, additional support will be needed for DMT in clinical contexts, with proper integration in place to minimise the impact of negative valence on mental health outcomes [40, 41]. Additionally, complementing our phenomenological findings, a recent study examined the impact of intravenous DMT under different doses in an extended-infusion protocol [42], where the psychedelic state can be maintained over longer periods of time. Our phenomenological results show some convergence with this study; we also found that higher doses led to increased retrospective ratings of entity encounters, visual hallucinations, and emotionally intense experiences. Reconstructing experiences through TET, however, has permitted a more direct mapping to the dynamic neural landscape induced through DMT.

Amongst the most strongly modulated dimensions, visual hallucinatory dimensions (i.e., Elementary Imagery and Complex Imagery) yielded the tightest associations with neural features, representing a broad increase in permutation entropy, reductions in oscillatory alpha power in tandem with broadband spectral shifts, and overall lower connectivity between gross EEG regions. The heightened intensity of these dimensions seem to broadly align with the fact that participants are more likely to enter a highly immersive, vivid and distinct state within the high dose. This begs the question: how could these neural features contribute to a state that so radically differs from normal waking consciousness?

Previous research has found that reductions in alpha oscillatory power are associated with visual hallucinations under psilocybin [43] and ayahuasca [13], a psychedelic brew containing DMT. Some theories of psychedelic action hypothesise that this reduction in alpha oscillatory power is related to its inhibitory function [44, 45], which may be a neurophysiological marker of the reduction of top-down predictions in a predictive processing framework of the brain [46]. The hypothesised neurobiological mechanism implicates 5-HT_2A_ agonism [43]. 5-HT_2A_ receptors are densely expressed in layer V pyramidal neurons, particularly within the primary visual cortex and transmodal cortical areas [10]. Activity within these cell-types have been found to produce oscillatory alpha activity [47], which may highlight how 5-HT_2A_ receptor agonism can induce a disinhibition of visual cortical activity, thereby facilitating the primarily visual hallucinatory DMT-state. However, as changes in oscillatory activity can arise from a multitude of biophysical factors [48–50], it is difficult to draw causal conclusions between specific neurobiological mechanisms and the radical shifts in conscious experience under DMT. Moreover, previous research has found that alpha power has demonstrable diagnostic utility to discriminate state in patients with disorders of consciousness [33, 51–53], with downregulation of alpha power, connectivity and graph theory measures, a strong neurophysiological hallmark of states of reduced consciousness. Conscious experience, while remarkably altered, remains preserved under the influence of DMT. Therefore, we highlight that oscillatory alpha power cannot be the sole neurophysiological driver of altered subjective experiences [31].

Permutation entropy yielded the second-strongest neurophenomenological associations amongst the neural features. Broadly, permutation entropy increases were associated with increases in TET dimensions (bar positive valence ones), indexing a shift towards more complex, rich and irregular neural signals. This broad association is in line with the entropic brain theory [19, 54], which posits that the entropy of the brain is associated with phenomenologically ‘rich’ states. This also concords with previous research, finding permutation entropy as a successful neural marker in discriminating conscious states under sleep [55–57], seizure [56, 58], and anaesthesia [59, 60]. However, interestingly, when computed with a tau value of eight, permutation entropy was inversely associated with phenomenological dimensions. As permutation entropy is computed through the ‘randomness’ of the probability distribution of signal motifs, a DMT-induced increase in high amplitude slow-wave activity, like [tau=8] permutation entropy indexes, may yield more predictable and consistent patterns, thereby reducing permutation entropy at this tau value. This exemplifies the importance of using more complex measures to parameterise neurophysiological entropy to more deeply understand the distinct spectral contributions [61], since in this study tau values of 1,2 and 4 show an inversion in the association to the dimensions of experience when compared to a tau of 8 (Figure 4).

LZ complexity, another information theoretic measure, yielded surprisingly few reliable neurophenomenological associations. This deviates from our most recent study showing that LZ complexity may be used to index TET-based dimensions of experience during breathwork, a non-pharmacological altered state of consciousness [31]. We argue that this discrepancy emerges due to both the relatively smaller sample used within this study^2^ (compared to [31]), and focusing our analyses on univariate dynamical associations. LZ complexity, as a relatively simplistic computational measure, may not be sensitive enough to capture subtle, fine-grained changes in experience and neural dynamics, and therefore may not be a suitable neurophenomenological marker in smaller scale studies [23, 62].

We emphasise the importance here of comparing and contrasting the relative utility of entropy-based neural markers in future research [63, 64].

Interestingly, by parameterising the PSD of the EEG signal we found minimal oscillatory delta power in the DMT condition. This conflicts with previous research highlighting an increase in delta power under the administration of DMT [6, 10] and ayahuasca [65, 66]. However, using traditional methodology, computing the relative power of the canonical frequency bands (See Supplementary Methods), we found that an increase in delta power was broadly associated with phenomenological dimensions within the DMT condition. These findings have demonstrable implications for cognitive neuroscience; parameterising the PSD, as done within this study, provides a more direct understanding of whether rhythmic or broadband neurophysiological changes are underpinning the phenomena of interest [36, 67]. We contest here that the increases in delta seen in previous research may be more reflective of broadband spectral changes, rather than increases in oscillatory delta components. Building on this, a recent landmark study of 5-MEO-DMT, another phenomenologically radical yet short-lasting psychedelic, not only found strong increases in slow-wave activity (<4Hz), but also characterised distinct spatiotemporal patterns of slow-wave activity compared to RS baseline [68]. More complex and dynamic methodologies may therefore be necessary to sufficiently study the vast-ranging effects of psychedelics on the brain, and to neurophysiologically taxonomise distinct conscious states and contents. We also encourage neurophenomenological approaches to strengthen the construct validity of future researchers’ findings.

We found that distinct families of neural markers had similar associations to univariate phenomenological dimensions. For example, Elementary Imagery, Temporality, and General Intensity were amongst the most strongly positively associated across all three marker families, with Bliss and Pleasantness yielding associations opposite to the thirteen other dimensions (e.g., increases in Pleasantness are associated with increases in oscillatory alpha power). This indicates two primary things: first, the top-performing dimensions may be most associated with the pharmacodynamics of DMT, phenomenologically typified by a complex visual hallucinatory state coupled with the dissolution of time. Therefore, these dimensions may be more tightly linked to between-subject effects, highlighting their strong association when summarising group-effect statistics, as in this study. Second, it seems that the neural markers are broadly indexing the same integrated phenomenological substrate, but with enough nuance to abandon the simple and single biomarker quest [69, 70]. It is not the case, for example, that connectivity markers are more associated with dissolutions of the self, whereas spectral changes are more associated with imagery-based dimensions. It may therefore be worthwhile to apply more multidimensional approaches to the simultaneous study of neural features to gain higher predictive power when examining both the brain and the mind. However, we acknowledge that such a multidimensional approach comes with interpretability challenges (as highlighted in [20]).

Regarding dose-response effects, neurophenomenological associations were broadly weaker in the medium dose compared to the high dose condition (Supplementary Figure S10). We propose that this may be due to comparatively weaker TET curves (e.g. Complex Imagery panel in Figure 2) that inevitably attenuate the association strength between experience dimensions and neural markers. For the evaluation of neurophenomenological relationships in psychedelic science, we argue that either stronger perturbations (such as the high dose, 40mg condition) or robust repeated-measures sample sizes [31] may be necessary to elucidate robust associations^3^.

While we present a novel approach to the study of experiential and neural dynamics, several limitations remain. First, it is difficult to estimate the exact doses ingested by each participant due to the method of combustion of DMT. The standardisation of dose-protocols, with verified purity and consistent administration procedures, is required to accurately and comprehensively characterise the effects of DMT across participants. Second, while our phenomenological results roughly converge with previous DMT research [9, 10, 42], our sample was predominantly male (17/19 participants) and hence, biased. Finally, a placebo condition was not applied within this research, and therefore placebo effects can not be ruled out. However, we highlight that this is a broad issue within psychedelic studies [71, 72], and may be more pertinent when assessing the clinical potential of psychedelic substances.

In conclusion, our results highlight how DMT impacted neural and experience dynamics in a dose-dependent manner. Using time-resolved measures of experience (TET) allowed the association of neural dynamics with phenomenological substates for every session, superseding the neurophenomenological potential of previous work with DMT and other psychedelics. It may be fruitful within the neuroscience of psychedelics, therefore, to implement such measures to more comprehensively map the commonalities and differences between distinct states of consciousness [73]. We also encourage researchers to continue to map the experiential and neural dynamics induced through psychedelics, and other altered states of consciousness [2], which can contribute to a more complete understanding of the relationship between the brain and the mind.

## Authorship contributions

Conceptualization: ELH, TB, ET, CP. Data analysis: ELH. Visualization: ELH, TB. Data Collection: ELH, CP, FC, TD, LDLF, DC, SM, NB, ET. Writing and editing: ELH, CP, FC, TD, LDLF, DC, SM, NB, ET, TB. Supervision: ET, TB.

## Acknowledgments

The authors wish to thank Eva Zaninetti for her assistance in creating the audio content for the sessions, Joaquim Streicher for creating the mobile application used to collect the TET data, and George Blackburne, Ravi Das, and Charlie Power for earlier discussions and development of the phenomenological dimensions.

## Methods

### Participants

The study was conducted in accordance with the Declaration of Helsinki and approved by the Committee for Research Ethics at the “Jose Maria Ramos Mejia” General Hospital (Buenos Aires, Argentina). Nineteen participants (17 males; 34.6±5 years; 4.7±5 previous DMT experiences) were recruited through word-of-mouth and social media advertisements. Screening for inclusion and exclusion criteria was conducted via a non-diagnostic interview with a mental health professional. During this initial conversation, participants were briefed on the experiment’s details, objectives, and eligibility criteria, and were provided with an electronic version of the informed consent form. Once eligibility was confirmed (criteria detailed below), participants were officially enrolled in the study and scheduled their dosing sessions, ensuring a minimum interval of two weeks between sessions.

### Inclusion and exclusion criteria

To participate in the study, individuals were required to have prior experience with ayahuasca or DMT (at least two sessions) and to abstain from psychoactive substances, including alcohol, caffeine, and tobacco, for at least 24 hours before participation. Eligible participants were between 21 and 65 years of age. Pregnant individuals were excluded. Participants who reported previous adverse psychedelic experiences resulting in lasting psychological difficulties or risky behaviours were also excluded.

A non-diagnostic mental health interview based on the Structured Clinical Interview for DSM-IV, Clinical Trials Version [74] (SCID-CT), was conducted in line with established guidelines [75]. Participants were excluded if they met DSM-IV criteria for schizophrenia or other psychotic disorders, bipolar disorder (types 1 or 2), or had first- or second-degree relatives with these conditions. Additional exclusions applied to individuals with substance abuse or dependence within the last five years (excluding nicotine), depressive disorders, recurrent depression, generalized anxiety disorder, obsessive-compulsive disorder, panic disorder, dysthymia, bulimia, anorexia, or any history of neurological disorders. Individuals scoring more than one standard deviation above the mean in trait anxiety, as measured by the State-Trait Anxiety Inventory [76] (STAI), or those on any form of psychiatric medication, were also deemed ineligible for participation.

### Experimental design

Within this study, we were able to employ a within-subjects blinded 2×2 design, with the factors dose (20mg and 40mg) and drug (DMT and RS) under naturalistic conditions. Doses were prepared and provided by an independent third-party facilitator. Neither the investigators nor the participants were informed about the specific doses administered, ensuring blinded conditions. The facilitator recorded this information and disclosed it to the research team only after the completion of the experimental phase, enabling the necessary data analysis while maintaining the study’s integrity. Prior to the first experimental session, participants were informed about the study details, with the definition of each phenomenological dimension explained (see Supplementary Methods for extensive dimension definitions). Participants were also provided with a mobile application (TET, Human Experience Dynamics, Ltd) which allowed them to practise conducting the experience tracing technique with a short online breath-based meditation session. Each of these practice traces were inspected by the primary researcher to ensure participants understood how to conduct the TET technique properly. In each dosing session, participants completed 10-minutes of RS baseline data with eyes closed, whilst EEG and physiological measures (ECG, GSR, accelerometer) were collected. Participants’ then traced the subjective intensity of the 15 phenomenological dimensions over the course of the 10minutes. Succeeding this, the participants smoked either 20mg or 40mg of DMT in its freebase form, through one singular and long inhalation. The order of the doses were randomised and counterbalanced. The participants’ then reclined with their eyes closed for the succeeding 20-minutes, whilst EEG and physiological recordings were taken. Throughout both the RS and DMT conditions, a series of bells were played at intervals of two-minutes, serving as temporal markers for the participants. Following the 20-minutes, participants’ were instructed to open their eyes. Once the participants’ expressed readiness, they traced the subjective intensity of each phenomenological dimension for the 20-minute session, with time-point zero representing the time in which the participant smoked the DMT. Once the participant had completed the traces, they briefly discussed their experience during the 20 minutes, in chronological order, with the audio digitally recorded, and EEG and physiological measures taken. Each participant completed two dosing sessions at least two weeks apart. Psychological support was available during each dosing session.

### Temporal experience tracing dimensions

To study subjective experience dynamics under the influence of DMT, we used Temporal Experience Tracing (TET) as our phenomenological methodology. TET has previously been used to study the phenomenological dynamics of different styles of meditation [29, 30], breathwork [31], chronic pain [32], conflict resolution [77, 78] and is currently being used to study the temporal dynamics of stress in autism [79]. With the methodology of TET, dimensions of subjective experience most relevant to the phenomenon of study are identified. These dimensions are sub-components of the contents of consciousness, relating to affective, attentional, or motivational aspects of experience. As mentioned above, participants complete the TET traces after both the RS or DMT conditions. That is, TET is a retrospective temporally dynamic quantitative phenomenological methodology.

During the session, bells were played to the participants at two-minute intervals, with one bell at two-minutes, two bells at four-minutes etc. These were also present in the TET graphs that participants had to complete as markers along the x-axis. Participants completed the TET graphs on a mobile application developed for a variety of other ongoing research studies using TET (e.g., [32]).

Fifteen phenomenological dimensions were used that loaded specifically onto the experience during DMT. To decide on these 15 dimensions, we examined the literature that investigated subjective experiences associated with DMT [4, 5, 7, 9, 10, 42, 80]. As mentioned above, participants received information about these 15 dimensions prior to the onset of this study. Participants received the definitions of each dimension in Spanish. Extensive definitions of each dimension can be found in the Supplementary Methods.

### Temporal experience tracing preprocessing

TET time series were imported in Javascript using Visual Studio Code Version 1.82.1. The data were preprocessed using a custom MATLAB script, where imported data were converted into a 15xN multidimensional matrix of subjective experience for each session and condition (RS or DMT), where N is the corresponding number of data points. Based on the assumptions of previous datasets [30, 31], each datapoint represented 30s of subjective experience, with N=20 for RS (10-minutes long) and N=40 for DMT (20-minutes long) conditions.

### EEG preprocessing

EEG data was collected with a BrainProducts 32-channel BrainAmp system using a sampling frequency of 250Hz. EEG data was preprocessed with EEGLAB using MATLAB. The EEG data was filtered between 0.5 and 45Hz. Noisy channels were rejected using an automatic procedure, where channels were rejected based on their kurtosis (threshold=5), and then manually scanned and rejected if identified as noisy (RS: x = 1.6 ±1; DMT: x = 2.2 ±1.1). For RS recordings, the first five seconds of the data were removed, and the data cut to a 10-minute recording time. For the DMT recordings, the data was cut to the point in which the participant exhaled after ingesting the DMT, and ended 20-minutes later. EEG data was re-referenced to the average signal, and split into five second epochs (1,250 samples). Epochs were rejected in a semiautomatic procedure, whereby epochs containing EEG amplitudes ±500µv or ±2SD of the amplitude of single channels and ±6SD of all channels were automatically rejected, and the data was manually scanned with any artefact laden epochs rejected (RS: x = 7.9 ±4.1; DMT: x = 24.3 ±15.2). ICA was applied separately to each condition per subject per session, and components related to non-neural artifactual elements such as eye movements and muscle artifacts were removed (RS: x = 1.9 ±1.1; DMT: x = 2.9 ±1.5). Following this preprocessing, two sessions from the DMT condition, and one from the RS condition were excluded from the EEG analysis due to over 50% of the epochs removed or high amplitude noise for the duration of the recording.

### Neural feature computation

We computed a variety of neural features broadly associated with the psychedelic-state. On the EEG data, we computed Lempel-Ziv complexity, permutation entropy, oscillatory power (delta, theta, alpha, beta), aperiodic spectral features (offset and exponent), weighted phase lag information, and weighted symbolic mutual information. Broadly, the neural markers computed fell into three families of markers: information theory, spectral, and connectivity. Spectral and information theory markers were summarised by computing the median average from each electrode in frontal (FP1, FP2, F3, F4, F7, F8, FZ, FC1, FC2, FC5, FC6), central (C3, C4, CZ), parietal (P3, P4, P7, P8, PZ, CP1, CP2, CP5, CP6), occipital (O1, O2), temporal (T7, T8, TP9, TP10, FT9, FT10), and global (all channels) regions. For connectivity measures, the median average was taken for pairings both within and between channels contained in the defined regions above (frontal, central, parietal, occipital, temporal). Global connectivity was computed as the median connectivity value for all channel pairings. Below is a summary of how to compute each neural marker, and our motivation for including each neural marker.

### Connectivity Measures

Previous research has found that weighted phase lag index (wPLI) and weighted symbolic mutual information (wSMI) may be useful markers to explore functional connectivity dynamics associated with conscious states [51, 81, 82]. In addition to this, previous research has shown that global functional connectivity increases as a function of psychedelic administration [10, 17, 83, 84], with a recent study demonstrating that a hyper-connected neural substate is associated with the rated intensity of Oceanic Boundlessness dimensions during a psilocybin session [85].

While both neural markers have been established as useful in the cognitive neuroscience of consciousness, wSMI and wPLI respectively focus on non-linear and linear interactions in the brain [86]. We therefore aimed to investigate how both of these interactions may be distinctly associated with phenomenology under the administration of DMT.

### Computation of wSMI

wSMI is a metric based on permutation entropy analysis [87]. For this metric, signals within each epoch are transformed into “symbols”, based on the ordering of the amplitude. Following this, mutual information can be computed between the symbolic transformation of pairs of electrodes (aka symbolic mutual information). However, based on the fact that EEG signals may be influenced by non-neural sources such as muscle artefacts or eye-blinks, a weighting system is introduced to weight identical or opposing values as 0, to prevent the conflation of non-neural signals with functional connectivity. We used k=3 as the parameter for symbol length, as previous research has used EEG systems with the same sampling frequency (250Hz) to analyse potential neural markers of conscious state [33, 34, 81].

The data was low-pass filtered with a formula of 80(Hz)/τ, with τ being a parameter referring to the temporal separation of each element within the symbolic transformation. wSMI was computed for four τ values [1, 2, 4, 8] between each electrode pair. For each five-second epoch, the wSMI value was averaged together for spatially distinct groupings of electrodes (frontal, temporal, central, occipital, and parietal). A global connectivity metric was also included, representing the average connection between all electrode pairs, thereby yielding 64 values per epoch. wSMI was computed using the nice toolbox [33]. It is noteworthy that wSMI can generate negative values [86]. These values were re-computed as 0, representing no information sharing between channels.

### Computation of wPLI

wPLI [88] is a metric based on the Phase Lag Index [89] (PLI). Both wPLI and PLI measure phase angle differences between two time series. However, while PLI discards phase differences at 0 or 180 degrees, which may be likely due to volume conduction, wPLI weights the magnitude of these phase differences, which prevents the risk of missing instantaneous interactions between electrodes [90, 91]. As above, wPLI was computed for each canonical frequency band (excluding gamma) between each electrode pair for the five-second epoch. The wPLI value was averaged together for spatially distinct groupings of electrodes (frontal, temporal, central, occipital, and parietal), with a global metric as an average of wPLI values between all channel pairings. As above, this leads to 64 wPLI values for each epoch.

### Spectral Measures

Previous research has found that serotonergic psychedelics are associated with broadband reductions in oscillatory power [15], with most prominent disinhibitory effects within the alpha frequency band [6, 9, 10, 12]. However, previous research has highlighted how aperiodic components (i.e., broadband/arrhythmic components) may be conflated with oscillatory activity [36, 67]. Aperiodic parameters (such as the aperiodic offset and exponent) have been related to the contents of consciousness in breathwork, a non-pharmacological altered state of consciousness [31]. Furthermore, other research has demonstrated that the aperiodic exponent can distinguish states of consciousness [92]. Aperiodic features are therefore becoming more integrated in neural analyses, increasingly being related to features of consciousness and cognition [36, 93, 94], further motivating our reason to study these aperiodic and oscillatory features in tandem.

### Computation of oscillatory components

We used the specparam (formerly fitting oscillations and oneover-f; fooof) methodology [36] to model both oscillatory and aperiodic components of the power spectra. For the oscillatory components, the specparam toolbox identifies peaks in the signal corresponding to oscillatory power, and models the offset and exponent of the PSD. We used the Welch method [95] on each five-second EEG epoch to compute the PSDs. We used 250-sample segments with 50% overlap (125 samples) and set the number of FFT points to 500, resulting in a frequency resolution of 0.5 Hz. Using specparam, we then parameterised each PSD to identify oscillatory peaks. To maximise the fit, we used [1, 8] as parameters for the peak width limits and 6 as the maximum number of peaks for each PSD. PSDs that did not yield sufficient fits (either the error value exceeding 0.125 or the r^2^ value below 0.9) were excluded from further analysis. We used the centre frequency parameter to determine whether the oscillatory component was localised within the canonical frequency bands Delta (δ: 1-4 Hz), Theta (θ: 4-8 Hz), Alpha (α: 8-13 Hz), Beta (β: 13-30 Hz), and used the “power” from the toolbox to parameterise the oscillatory amplitude. If multiple peaks were identified in the same frequency band and the same PSD, the “power” parameter was summated. If no peaks were identified, the power value was 0 by default. Gamma activity was excluded from this analysis as it may more likely reflect muscular activity in EEG [96]. When computed and averaged for each spatial region (frontal, temporal, central, parietal, occipital, and global) a total of six values per frequency band per epoch were generated.

### Computation of aperiodic components

As above, the specparam toolbox was used to parameterise the PSDs for each EEG epoch for each channel. The offset, reflecting broadband shifts in power, and the exponent, reflecting the rate of change over the course of the PSD, were the aperiodic parameters that were output. PSDs that did not yield sufficient fits (either the error value exceeding 0.125 or the r^2^ value below 0.9) were excluded from further analysis. When computed and averaged for each spatial region (frontal, temporal, central, parietal, occipital, and global), a total of 12 aperiodic values per epoch were generated.

### Information theory measures

Due to previous theories highlighting that the entropy of the brain is related to phenomenology [18, 19], we also computed information theory measures on the EEG data. The methods broadly purport to measure the (un)predictably or selfsimilarity of the EEG signal, through nonlinear approaches, thus moving away from the reliance on analysing oscillatory components.

### Lempel-Ziv complexity

Lempel-Ziv (LZ) complexity is a relatively simple method that has been robustly applied to cognitive neuroscientific work on consciousness [20, 21, 24, 97, 98]. LZ complexity is computed through the binarisation of the signal around its median, where all values above the median are 1, and all values below the median are 0. The binarised signal is then algorithmically scanned sequentially for novel patterns within the signal [99], creating a ‘dictionary’ of binary sequences for each timeseries. The length of this dictionary is a standardised measure of the complexity of the signal.

We computed LZ complexity from channels within gross EEG regions (frontal, parietal, central, temporal, and occipital) in two distinct ways. In the first form, timeseries from all channels within one spatial area and epoch were concatenated together, with the computations above applied to the single concatenated signal (concatenated LZ; LZc). LZ was then normalised by dividing the raw LZ value by the LZ value computed through the same binary input that is randomly shuffled [24]. Normalising on this basis provides a complexity measure falling within 0 to 1. In the second form, the LZ computation above was applied to each channel separately, with the results averaged (summed LZ; LZsum). This generated two measures of complexity for one of the five spatial areas, and one global metric, leading to 12 LZ complexity values per epoch.

### Permutation entropy

Permutation entropy [87] is a method to determine the complexity of timeseries that is robust to low signal-to-noise ratios [33]. For permutation entropy the signal is transformed into groups of symbols (length = n), separated by τ samples. In this regard, the parameter τ represents the frequency of interest, corresponding to either theta [τ = 8], alpha [τ = 4], beta [τ = 2], or gamma [τ = 1] frequencies. Shannon’s entropy is then computed on the probability distributions of the symbols over the timeseries. We calculated permutation entropy with four τ parameters [1, 2, 4, 8], and n = 3. The latter is used as previous research has found that this is a robust parameter to use when sampling at 250Hz and investigating markers of consciousness [33, 34]. When computing for four τ parameters, five spatial regions and one global measure, 24 values per epoch were generated.

### Temporal alignment of neurophenomenological data

Due to the distinct preprocessing of the TET data (with 30s ‘phenomenological epochs’) and the EEG data (five-second epochs), each neural feature was averaged over six epochs to temporally align the neural and phenomenological data. Succeeding this, any data that was above three standard deviations of the mean were transformed to nans and then replaced by the average of the two neighbouring neural feature datapoints within that session (i.e., within-session imputation).

### Hypotheses

For the phenomenological analyses, we hypothesised that the 40mg dose condition would be significantly higher in intensities of all phenomenological dimensions, with most pronounced differences in General Intensity, Elementary Imagery, and Complex Imagery. Further, we also hypothesised that the 40mg dose will be significantly different to the average RS baseline for longer periods of time in all dimensions when compared to the 20mg dose.

For the neurophenomenological analyses, we hypothesised that information theory markers, as well as oscillatory alpha and delta power, would be most strongly associated with phenomenology. This hypothesis is based on previous neuroscientific research with DMT [6, 9, 10], findings implicating complexity measures as robust markers of conscious state [64], and theories of psychedelic action [18, 19]. Additionally, we hypothesised that non-linear connectivity markers, (i.e., weighted symbolic mutual information; wSMI), would surpass linear connectivity markers (i.e., weighted phase lag information; wPLI) in neurophenomenological associations, supported by previous research in bistable perception paradigms [100, 101], and deep meditative states [102].

### Statistical analysis

For the phenomenological analysis, we computed linear mixed models to evaluate (a) the phenomenological differences between 40mg and 20mg DMT for each experience dimension, and (b) the difference between RS conditions and DMT conditions for each phenomenological timestep, with dose as an interaction effect. The advantage of using linear mixed models is that participants and sessions can be factored in as random effects, and we did not need to exclude participants that had TET or EEG data missing from the analysis, therefore maximising our statistical power within this more complex repeated-measures design.

For the neurophenomenological analysis, we computed linear mixed models to evaluate the association between each neural feature (194 in total within the primary analysis) and each phenomenological dimension. In total, 2910 linear mixed models were computed (450 for information theory measures, 540 for spectral measures, and 1920 for connectivity measures).

For all analyses, FDR multiple comparisons [103] were computed to adjust the p-values of the models using the multicmp function in MATLAB [104]. Further, all continuous data were transformed to have mean of 0 and variance of 1, thereby providing a comparable estimate of effect size across all neural features and phenomenological dimensions [35].

---

**Figure S1:**
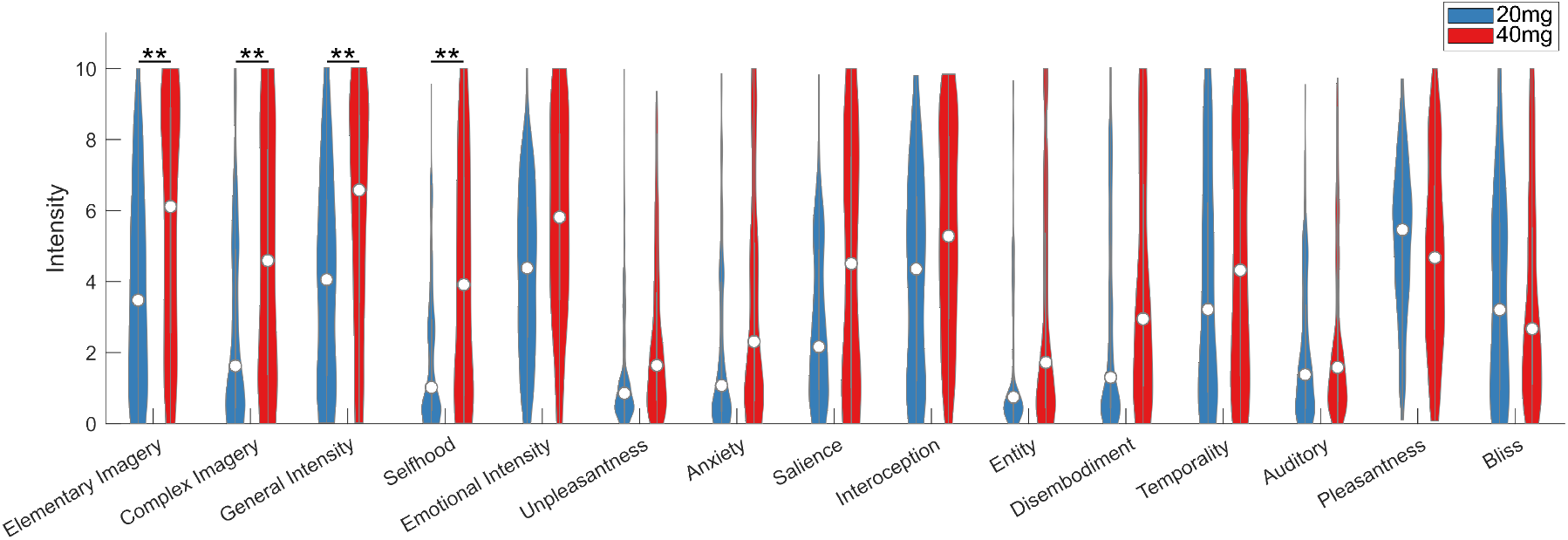
Distributions of phenomenological TET data for all participants and both DMT doses. Violin plots of the distribution of the TET data across all participants, doses, and phenomenological dimensions. When statistically analysing the difference between dimension intensities between doses, however not factoring time in to the linear mixed models, only Elementary Imagery, Complex Imagery, General Intensity and Selfhood were significantly different after FDR correction. This emphasises the necessity of factoring time into phenomenological models to gain a more nuanced understanding of the temporal dynamics of experience within the DMT-state. \*\**p*<0.01

**Figure S2:**
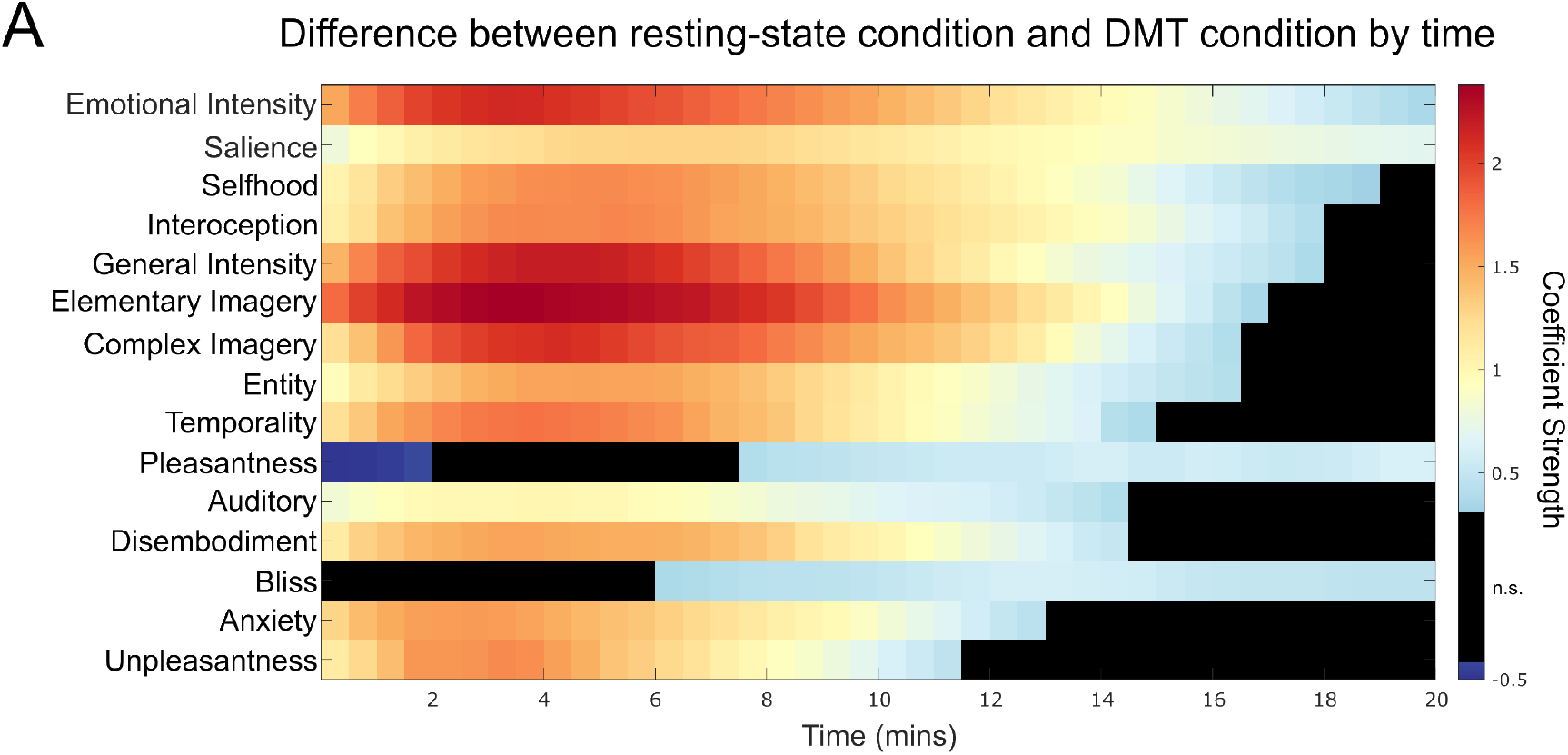
Time-resolved differences between resting-state phenomenology and DMT phenomenology regardless of dose. Each timestep presents a linear mixed model coefficient indicating whether that TET datapoint in time was significantly different to the resting-state average for that session. An exemplary mixed model to compute this was ‘Emotional Intensity ∼ Time*Dose + (1| Subject:Session)’. The main effects here are the coefficient strengths. As each phenomenological dimension was standardised to mean = 0 and variance = 1, the coefficients are comparable across phenomenological dimension, and therefore an indication of the strength of the effect (i.e., effect size). Phenomenological dimensions on the y-axis are ordered based on the overall strength of effects over the whole 20-minute session.

**Figure S3:**
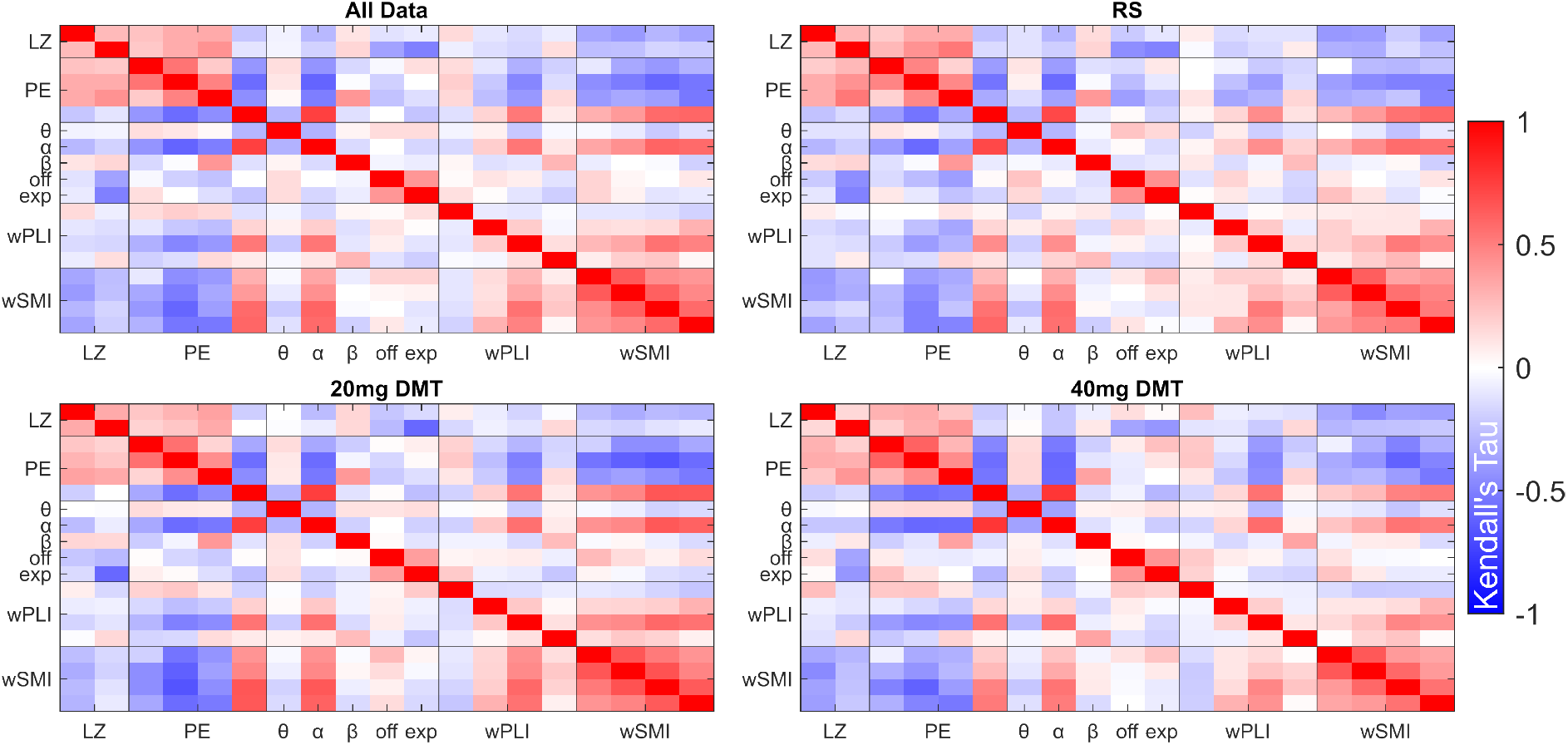
Covariance matrices between each global neural feature across different conditions highlight consistent relationships between neural features. Covariance matrices were computed using Kendall’s Tau correlation between each neural feature (averaged from all 31 channels) across all of the data (top left), the resting-state condition (top right), the 20mg DMT condition (bottom left), and the 40mg condition (bottom right). Across conditions, there are consistent relationships across the neural features.

**Figure S4:**
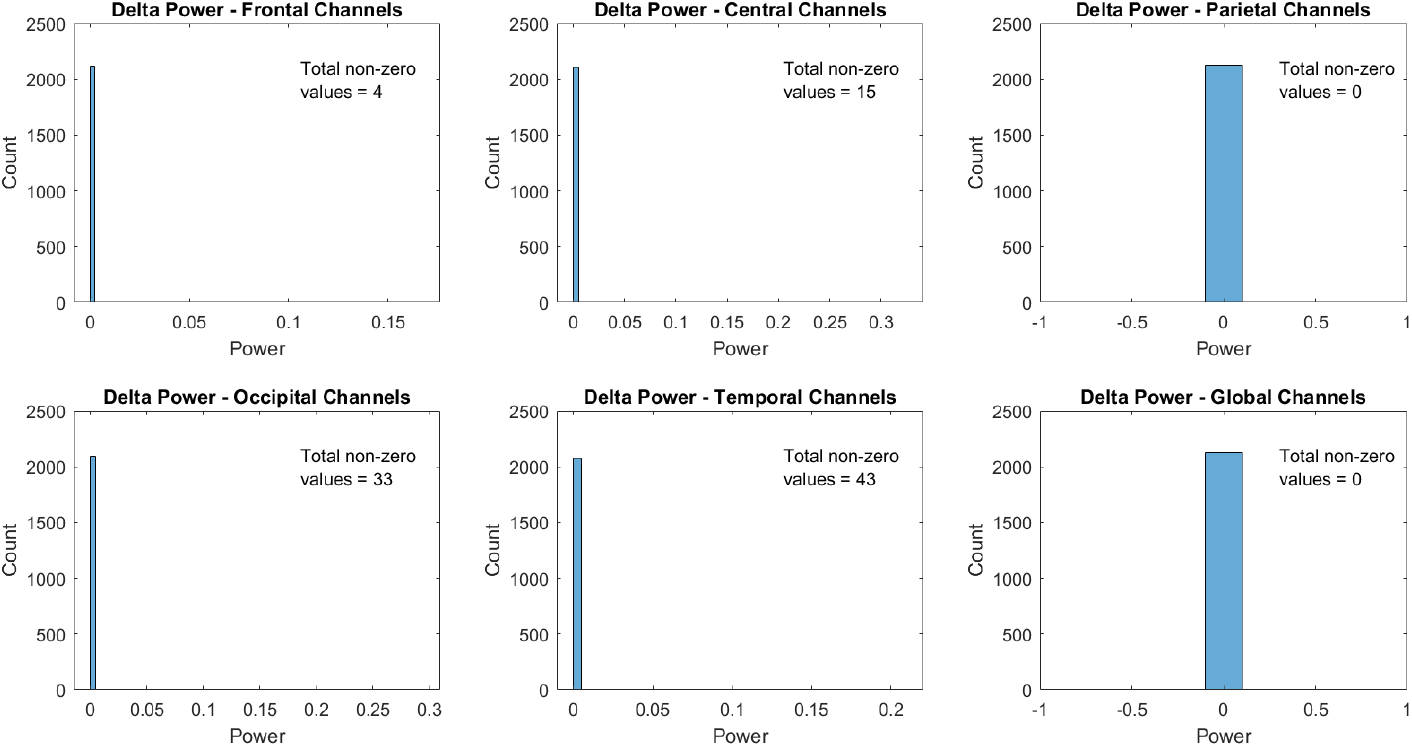
Little delta oscillatory power when computing through the specparam methodology. The distribution of oscillatory delta power when computed through the parameterisation of the power spectral density (PSD) using the specparam methodology [36]. Each histogram presents the distribution of oscillatory delta power for the respective channel groupings. Even for the temporal channels, which had the highest amount of non-zero power values, this only represented under 2% of the data. Due to this, oscillatory delta power was excluded from the analysis.

**Figure S5:**
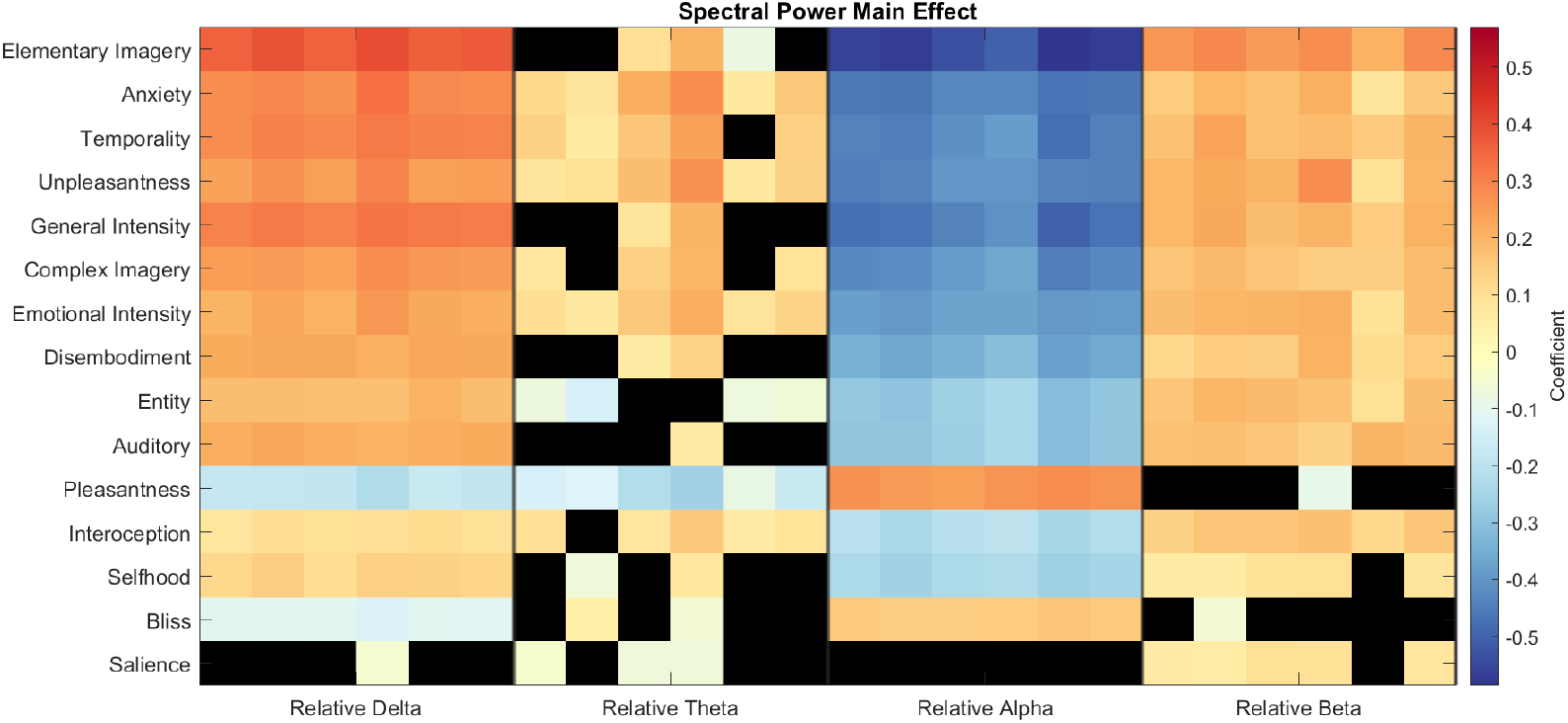
Neurophenomenological associations for relative power of PSD for the canonical frequency bands. The coefficient matrix presents the main effect coefficients for linear mixed models computed with relative power in the delta (1-4Hz), theta (4-8Hz), alpha (8-13Hz), and beta (13-30Hz) frequency bands. Relative power was estimated as the percentage of total power within the frequency band over the whole PSD within the 1-30Hz range. Gamma is therefore excluded from this analysis (results with gamma are provided below in Fig. S6). Within the figure, black values are non-significant after FDR correction. Focusing on relative delta power, we found broadly positive and strong neurophenomenological associations. Phenomenological dimensions on the y-axis in order of strength of association across all spectral neural markers.

**Figure S6:**
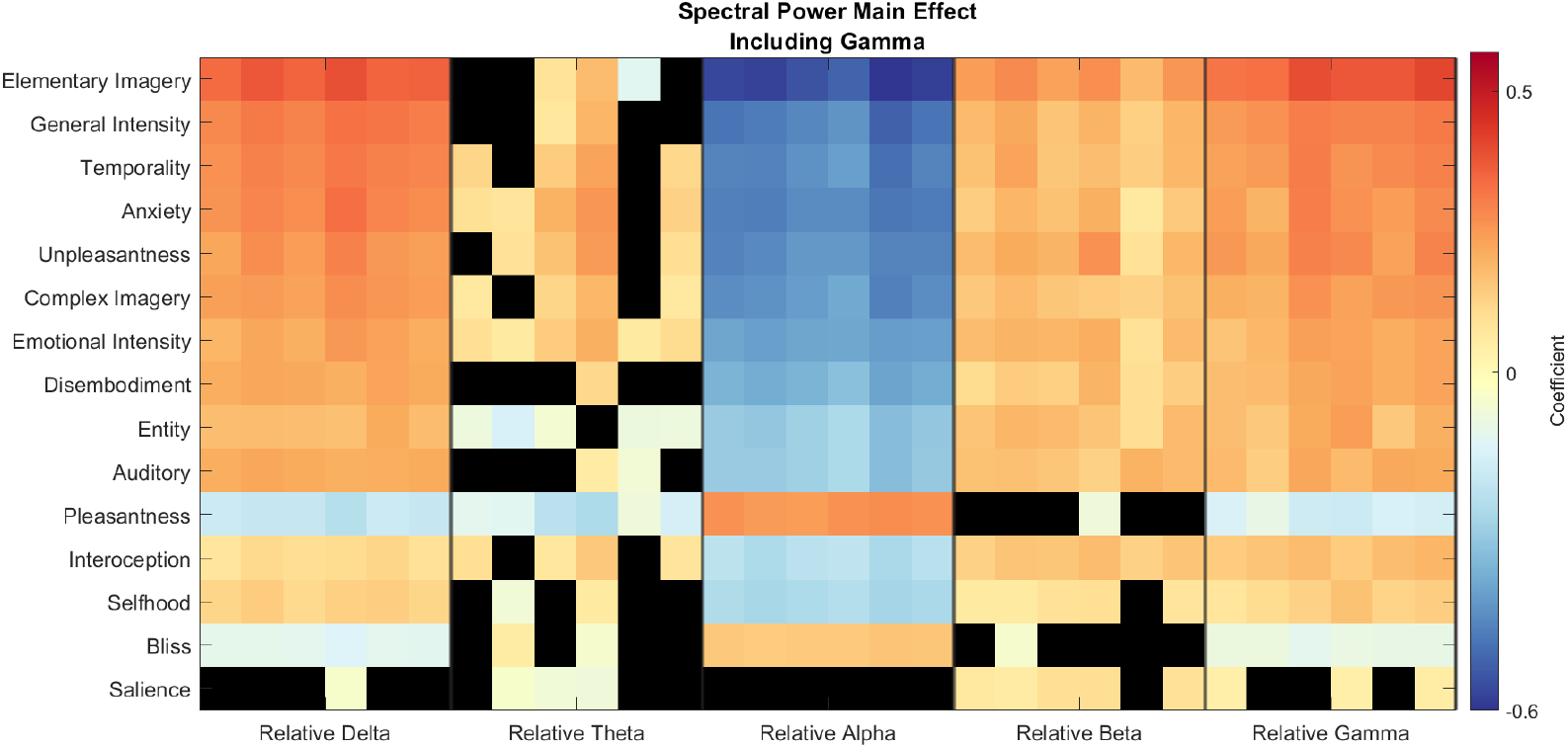
Neurophenomenological associations for relative power of PSD for the canonical frequency bands, gamma power included. As in Fig S3, the coefficient matrix presents main effects for linear mixed models computed between each neural feature and each phenomenological dimension. Unlike Fig. S3, however, the relative power in each frequency band is computed from PSDs in the 1-45Hz frequency band, which therefore includes relative gamma power. Black values are non significant (*p*>0.05) after FDR correction.

**Figure S7:**
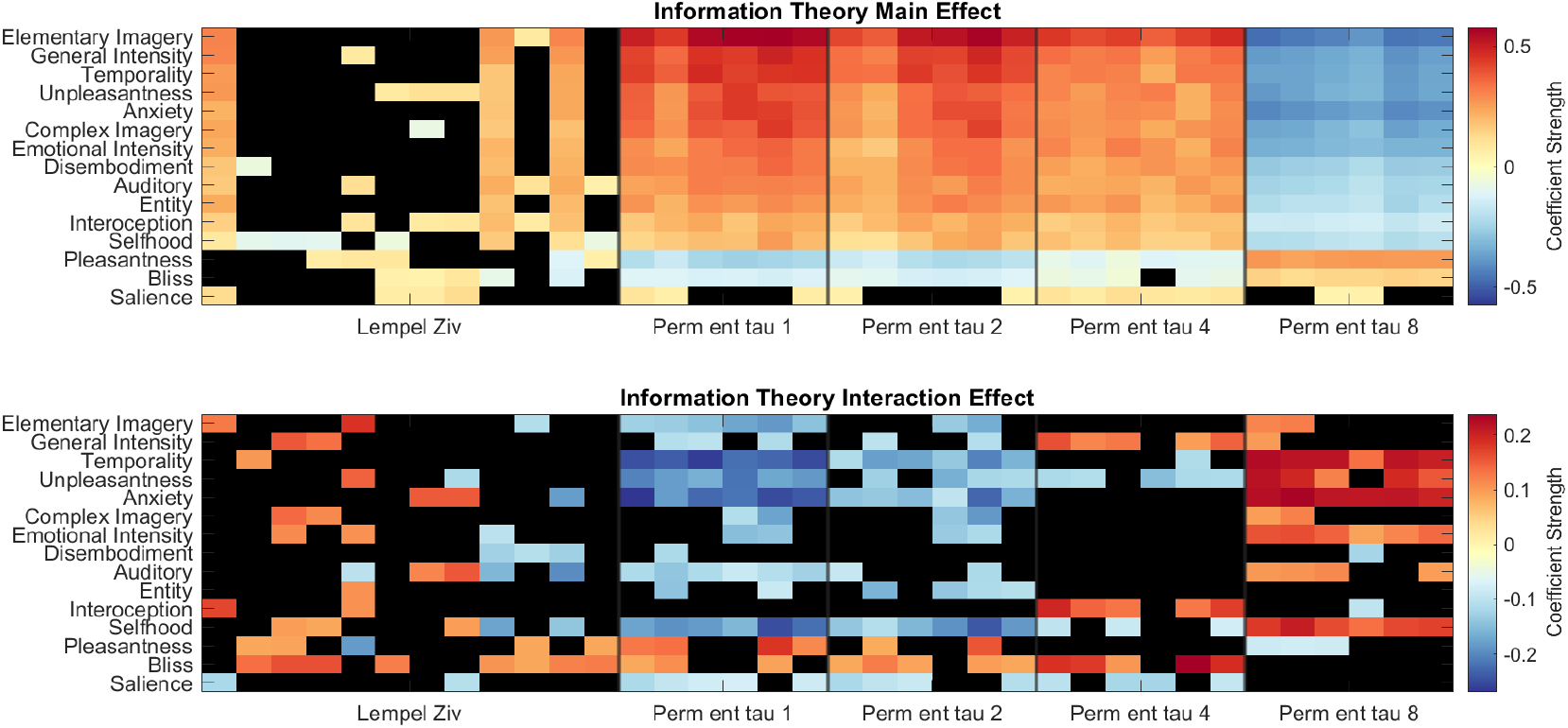
20mg DMT dose generally yields weaker neurophenomenological associations for information theory measures. Main (top) and dose interaction (bottom) effects for the neurophenomenological associations between phenomenological dimensions and information theory neural markers. The dose interaction effects highlight that there is a difference in association strength when comparing medium dose to high dose. For example, the 20mg DMT dose had significantly weaker association between permutation entropy (τ = 1) and Elementary Imagery. Black values are non significant (*p*>0.05) after FDR correction.

**Figure S8:**
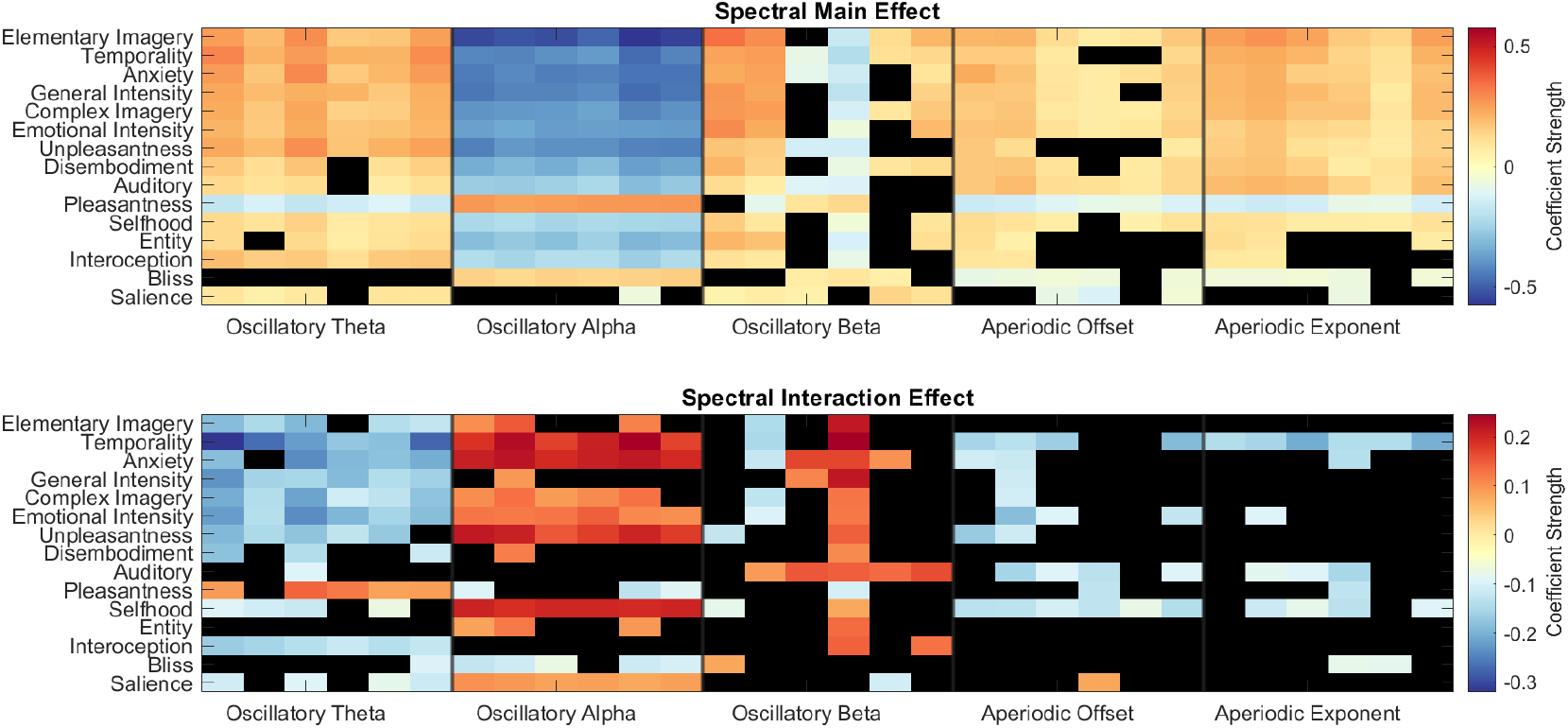
20mg DMT dose generally yields weaker neurophenomenological associations for spectral measures. Main (top) and dose interaction (bottom) effects for the neurophenomenological associations between phenomenological dimensions and spectral neural markers. The dose interaction effects highlight that there is a difference in association strength when comparing low dose to high dose. For example, the medium dose DMT condition had significantly weaker associations between oscillatory alpha power and Selfhood, as the interaction coefficient strength is positive, whereas the main effect is negative. Black values are non significant (*p*>0.05) after FDR correction.

**Figure S9:**
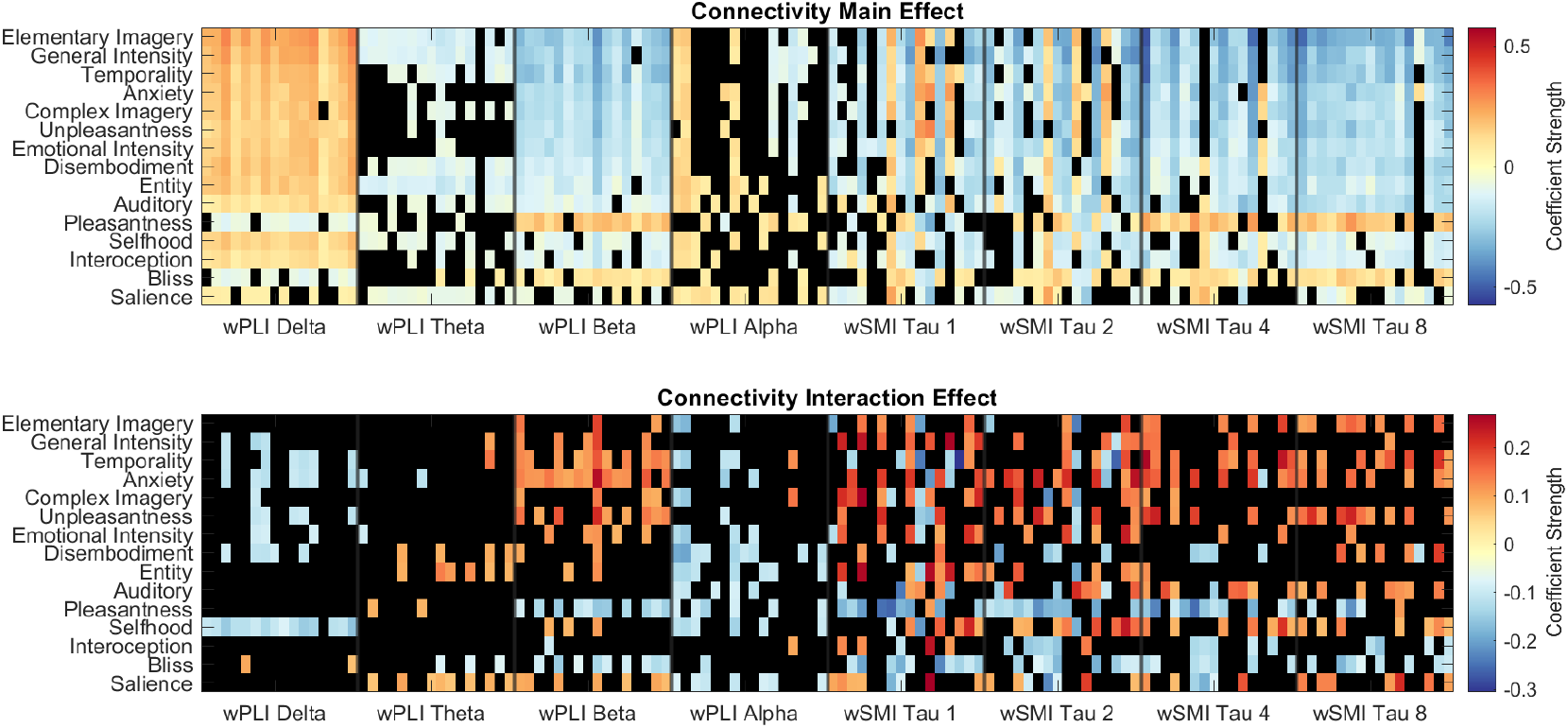
20mg DMT dose generally yields weaker neurophenomenological associations for connectivity measures. Main (top) and dose interaction (bottom) effects for the neurophenomenological associations between phenomenological dimensions and connectivity neural markers. The dose interaction effects highlight that there is a difference in association strength when comparing medium dose to high dose. For example, the medium dose DMT condition had significantly weaker associations between Beta wPLI and Anxiety, as the interaction coefficient strength is positive, whereas the main effect is negative. Black values are non significant (*p*>0.05) after FDR correction.

**Figure S10:**
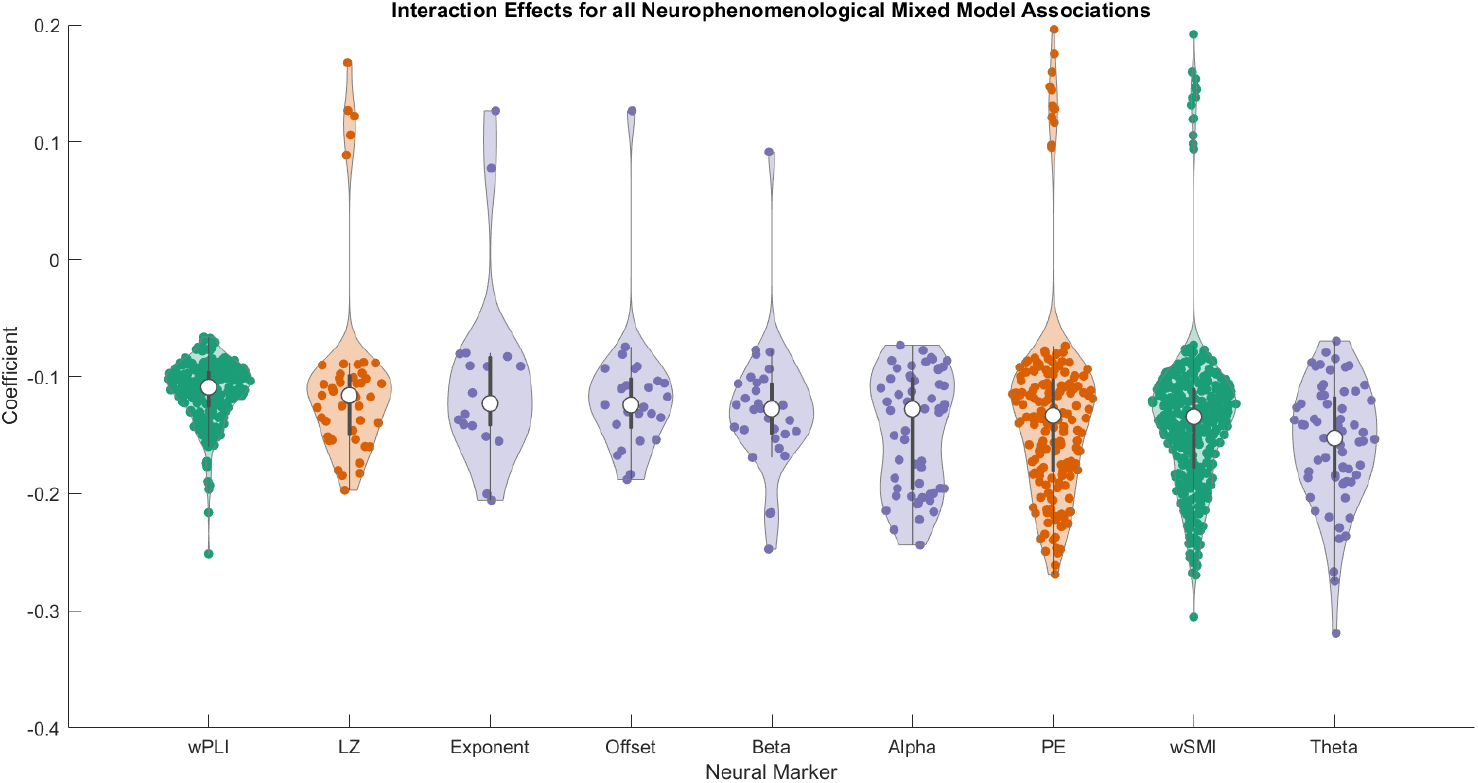
20mg DMT dose mitigates neurophenomenological associations across neural markers. The distribution of significant interaction effects (after FDR correction) across all neural markers. The sign of the coefficients were adapted to represent whether the 20mg dose increased or decreased the main effect, whereby negative coefficients represent a reduction in neurophenomenological association, and positive coefficients represent an increase in neurophenomenological association. To do this, if the interaction effect and main effect had the same sign (both positive or negative), then the interaction effect was transformed to positive. However, if they were different signs, the interaction effect was transformed to negative. Doing so makes the interaction coefficients comparable across all neural markers, regardless of whether they had negative or positive associations with phenomenology. The majority of the interaction effects were below 0, indicating that the 20mg dose usually reduced neurophenomenological associations across all neural markers. Neural markers are ordered in terms of their median strength (shaded in white in the centre of each violin plot).

**Figure S11:**
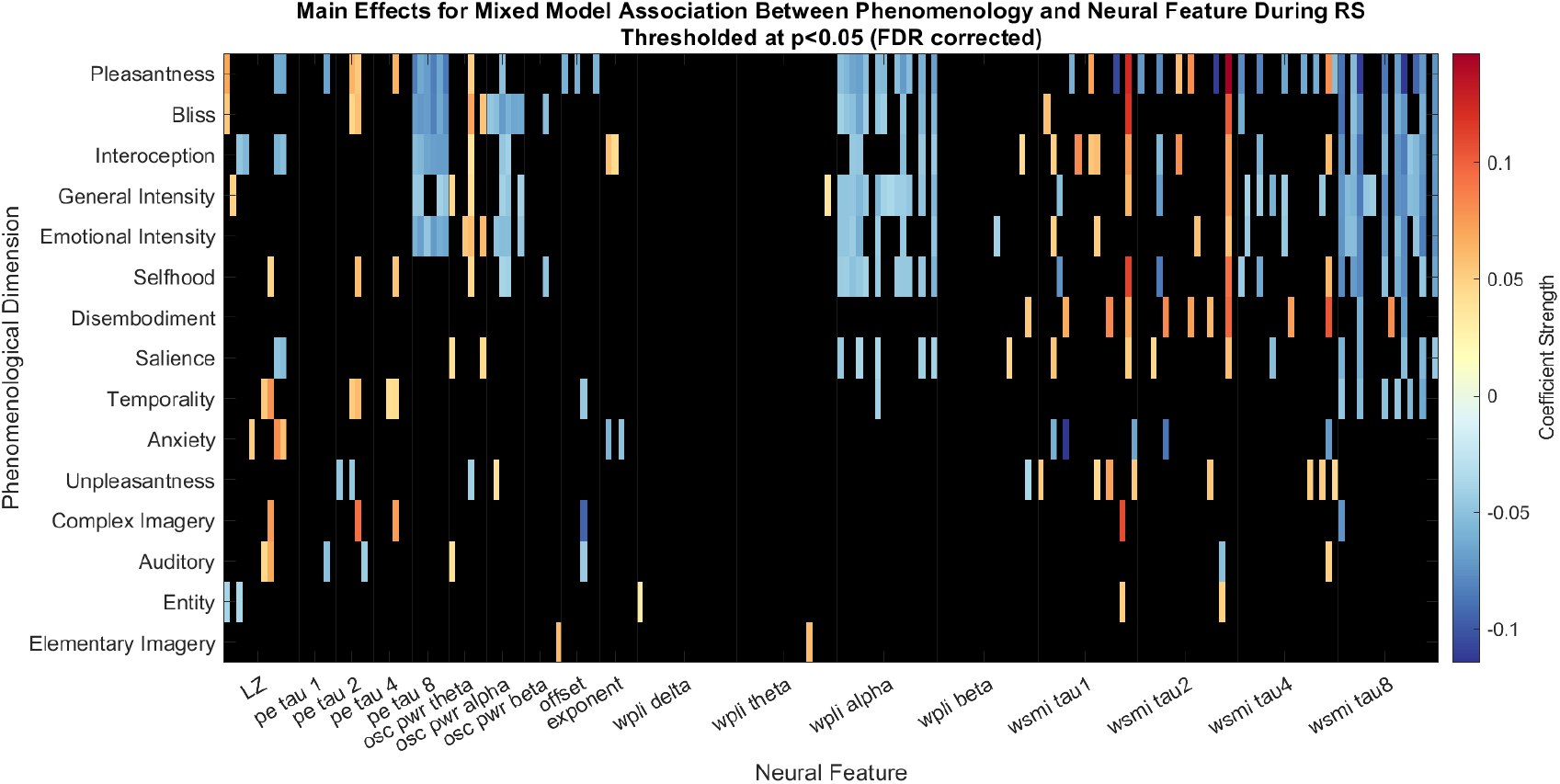
Little neurophenomenological associations across all neural markers and phenomenological dimensions in the resting-state condition. This coefficient matrix is the output of the mixed models examining the association between each neural marker and phenomenological dimension for the resting-state data. An exemplary linear mixed model is ‘Pleasantness ∼ global alpha power + (1 | Subject:Session)’. Dose was not included as an interaction effect (as in the primary analyses within the paper), due to no expectation of session-related differences. Black values within the matrix did not reach significance of *p*>0.05 after FDR correction.

## Supplementary Methods

### Phenomenological dimension definitions

Below is an elaboration of the definitions of each phenomenological TET dimension used within the study.

### Pleasantness

This dimension is specifically concerned with the felt intensity of pleasantness for experiences over time. Pleasantness refers to the subjectively felt ‘goodness’ to the experience at that moment in time. Low intensity ratings on this dimension does not mean that an experience is unpleasant, it simply means that it does not feel intrinsically pleasant at that time. Positive mood states are associated with a sub-component of the mystical experience questionnaire [105], highly related to the psychedelic state [106]. We included this dimension to probe the overall valence of the experience.

### Unpleasantness

This dimension is specifically concerned with the intensity of unpleasantness for your experiences over time. Unpleasantness refers to the subjectively felt ‘badness’ to the experience at that moment in time. Similar to the above, low intensity ratings on this dimension does not mean that an experience is pleasant, but simply means that it does not feel intrinsically unpleasant at that time. We included this dimension as some aspects of the psychedelic experience can be challenging [107], and we therefore wanted to probe the overall valence of the experience.

### Emotional Intensity

This dimension is specifically concerned with the intensity of emotions, rather than the valence. The psychedelic experience can heighten emotional intensity [108], and can also lead to experiences of catharsis or emotional breakthrough [109], which can be highly emotionally intense.

### Elementary Imagery

Elementary imagery is specifically concerned with low-level visual sensations, such as flashes, colours and geometric patterns. Elementary imagery is a sub-component of the 11 Dimensions of Altered States of Consciousness questionnaire [110] (11D-ASC), and robustly increases as a function of DMT administration [6, 9]. Higher ratings on this dimension refer to a predominance of elementary imagery within conscious experience.

### Complex Imagery

This dimension is also concerned with sensations in the visual domain, however specifically relates to visual sensations that are more complex in their nature. For example, experiencing vivid scenes, pictures of the past, or fantastical visions. As above, Complex Imagery is a sub-component of the 11D-ASC, and significantly increases under the administration of DMT [6, 9]. Higher intensity ratings refer to the fact that complex imagery is highly predominant within conscious experience.

### Auditory

This dimension is specifically concerned with sensations in the auditory domain, whether that be more awareness of sounds from the external world or the hallucinatory experience of sounds. Higher intensity ratings on this dimension mean that there is a large predominance of auditory perception at that time. Low intensity ratings means that there is a low predominance of auditory perception at that time. Recent research found significant auditory alterations at higher doses of DMT [42].

### Interoception

This dimension refers to the intensity of sensations within the body. High ratings in the dimension were associated with strong somatic sensations like pulsing, intense heaviness, or the sensation of ‘energy’ within the body. This is sometimes referred to as ‘body load’. Low ratings on this dimension were associated with an unawareness or unsensational occurrence of feelings within the body. Previous research has found that the intensity of bodily sensations dynamically increases as a function of DMT dose [42].

### Bliss

This dimension refers to the experience of bliss or profound peace. This is an inherently positive experience, defined by boundless pleasure. Bliss is a sub-component of the 11D-ASC [110]. Previous neurophenomenological research with breathwork has found that the retrospectively rated intensity of bliss during breathwork as an altered state of consciousness is associated with neural Lempel-Ziv complexity [31]. Subjective experiences cultivated by DMT are also associated with an increase in blissful state [6, 9, 42].

### Anxiety

This specific dimension relates to the experience of dysphoria, an antonym of euphoria. As mentioned above, psychedelics can cultivate challenging experiences [107], with DMT inducing transient dose-dependent increases in anxiety [42]. Low intensity ratings on this dimension did not signify a euphoric experience, just that there was little to no anxiety in subjective experience.

### Entity

Previous research has shown that the DMT experience may induce the felt presence of ‘autonomous entities’ [11, 111, 112]. These entities are commonly described as guides, spirits, aliens, and helpers [111], and are experienced as having their own sentience. The intensity of this dimension relates to the presence of entities in subjective experience, with low ratings meaning no experience of entities.

### Selfhood

Previous research has sought to investigate how selfhood dissolves as a function of psychedelic administration, as demonstrated by the development of the ego dissolution inventory [113] (EDI). Previous research has found that EDI scores were significantly higher under DMT compared to placebo [114], indicating a dissolution in selfhood in this state. High intensity ratings on this dimension represented high deviations in the experience of the self, whereas low intensity ratings on this dimension represented a typical experiencing of the ‘self’, typified in normal waking consciousness.

### Disembodiment/Embodiment

Disembodiment refers to the experience of not identifying with one’s own body [115]. The antonym of this, the feeling of identifying with and normally perceiving one’s body, has frequently been investigated and modulated in studies on self-consciousness [116]. Disembodiment forms a specific sub-component of the 11D-ASC [110], and has been modulated through the administration of psychedelics [117–119]. In particular, micro-phenomenological interviews with participants revealed that bodily alterations were one of three broad dimensions constituting the DMT experience [9].

### Salience (Meaning)

The dimension of salience refers to a subjective sense of the moment being deeply significant and important. Salience was included as serotonergic psychedelics have been found to elicit extraordinarily significant and meaningful subjective experiences [120–122]. High ratings of this dimension refers to intensely meaningful moments of experience, whereas low ratings refer to moments that did not feel especially meaningful, significant, or important.

### Temporality

This dimension refers to the subjective experience of time. During normal waking consciousness, time may be experienced in a linear and smooth fashion, however, under the influence of DMT, time may lose its normal structure and shape. The transcendence of time (and space) forms a subcomponent of the mystical experience questionnaire [105] (MEQ), commonly used to assess subjective experiences of psychedelics. Further, increases in this subcomponent of the MEQ after smoking DMT was associated with increases in gamma power [6]. High ratings on this dimension are related to how intensely the sense of time felt disrupted.

### General Intensity

This dimension refers to the general subjective intensity of the effects of DMT. Previous research has found neurophenomenological associations with neural Lempel-Ziv complexity and the subjective intensity of DMT [9] and other psychedelics [21].

### Computation of relative power for supplementary analyses

In addition to the primary neural feature computation, we also computed spectral power in supplementary analyses in a more traditional approach (results presented in Figs S3 & S4). The power in each frequency band was computed as the integral of the PSD at the corresponding frequency values. From this, we present “relative power”, which is computed through the division of the power in each band by the total power, providing a percentage of total power that each frequency band covers. This mitigates issues related to inter-individual variance due to, for example, eye movements and electrode impedance. This was computed both with and without gamma power.

## Footnotes

1 For the median and standard deviation of the coefficients, we converted all negative values into non-negative, to make the coefficients comparable across all neural features (e.g., permutation entropy can be compared to oscillatory alpha power despite the former having positive neurophenomenological associations and the latter having negative associations). Further, we included non significant coefficients.

2 It is noted, however, that despite the relatively small absolute sample size within this study, the dynamic analyses applied to both EEG and experience data has strengthened its statistical power.

3 However, we acknowledge the difficulty of obtaining larger sample sizes with multiple dosing sessions when working with psychedelic substances.

